# Prime assembly with linear DNA donors enables large genomic insertions

**DOI:** 10.1101/2025.06.16.659978

**Authors:** Bin Liu, Andrew Petti, Xuntao Zhou, Haoyang Cheng, Lin Zhou, Tingting Jiang, Erik J. Sontheimer, Wen Xue

**Affiliations:** RNA Therapeutics Institute, University of Massachusetts Chan Medical School, Worcester, MA, USA; Department of Molecular, Cell and Cancer Biology, University of Massachusetts Chan Medical School, Worcester, MA, USA; Icahn Genomics Institute, Department of Immunology and Immunotherapy, Department of Oncological Sciences, Department of Genetics and Genomic Sciences, Icahn School of Medicine at Mount Sinai, New York, NY 10029, USA; Li Weibo Institute for Rare Diseases Research, University of Massachusetts Chan Medical School, Worcester, MA, USA; Department of Molecular Medicine, University of Massachusetts Chan Medical School, Worcester, Massachusetts, USA; Department of Biochemistry and Molecular Biotechnology, University of Massachusetts Chan Medical School, Worcester, MA 01605, USA

## Abstract

Targeted insertion of large DNA fragments has promising applications for genome engineering and gene therapy^1,2^. Twin prime editing (PE) guide RNAs (pegRNAs) have enabled relatively large insertions, but the efficiency remains low for insertions greater than 400 base pairs^3–6^. Here we describe a Prime Assembly (PA) approach for the insertion of large DNA donor fragments, whose ends are designed to overlap with the flaps generated by twinPE. We used PA to insert one, two, or three overlapping DNA fragments, with total insertion sizes ranging from 0.1 to 11 kilobase pairs. An inhibitor of non-homologous end joining (NHEJ) enhanced both the efficiency and precision of insertions. PA relies on DNA templates that are easily produced and does not require co-delivery of exogenous DNA-dependent DNA polymerases. Our study demonstrates that PA can initiate “Gibson-like” assembly in cells to generate gene insertions without double-stranded DNA breaks or recombinases.

## Introduction

Many genetic diseases are caused by diverse mutations across various loci, necessitating precise therapeutic strategies^7^. However, current genome editing approaches struggle to efficiently or accurately insert or replace large DNA fragments. Traditional approaches for targeted integration of gene-sized DNA fragments rely on double-strand break (DSB) repair via non-homologous end joining (NHEJ) or homology-directed repair (HDR)^8^. Unfortunately, these methods have significant limitations. DSBs can cause uncontrolled insertions or deletions (indels) and activate p53, resulting in DNA damage and cytotoxicity^9,10^. In the presence of a donor DNA, a DSB can be precisely repaired by HDR, but HDR is usually inefficient for large DNA insertions in primary cells and *in vivo* contexts^11–13^. Thus, developing methods for precise, efficient large DNA integration remains a key challenge in genome editing^14^.

Recent advances in prime editing (PE)-associated technologies, combined with serine integrases, have enabled large insertions into the genome^3,15–17^. However, these approaches often produce junctional scars at insertion sites, such as *attP*-*attB* junctions or recombination-induced misalignments, leading to small insertions, deletions, or substitutions. These imperfect editing events could have adverse functional consequences^3,16,18^. Moreover, the *in vivo* or clinical safety of integrases remains poorly characterized, and the substantial size and structural complexity— including PE/recombinase effectors, plasmid donors, and pegRNA templates—pose considerable challenges for efficient delivery. These factors present significant translational barriers, although various studies have aimed to address these and other limitations^3,17,19–26^. Current PE-based systems, such as PASSIGE and PASTE^3,16^, primarily use plasmids as donor templates. However, plasmids exhibit poor uptake *in vivo* and can elicit innate immune responses due in part to unmethylated CpG motifs, leading to inflammation^27^.

Given these challenges, there is an urgent need for a scarless, plasmid-free, and integrase-free system capable of achieving efficient, precise large DNA insertions, particularly for therapeutic applications.

## Results

We and others have developed twinPE, DNA polymerase editing (DPE), and related technologies^3,5,33,16,17,20,28–32^. However, these approaches are not yet efficient at achieving gene-sized insertions (>800 bp), likely due to multiple factors including pegRNA instability, limited DNA synthesis activity of the exogenous reverse transcriptase (RT) or DNA-dependent DNA polymerase, DNA repair bottlenecks, and other constraints. Because prime editors efficiently generate 30- to 40-nucleotide (nt) flaps, we hypothesized that a linear double-stranded DNA (dsDNA) donor, matching the flaps generated by twinPE, could bypass the limitations of exogenous polymerase-mediated synthesis and enable large DNA insertions.

To test this, we developed a Prime Assembly (PA) approach, in which the ends of a PCR-produced dsDNA donor are homologous to programmable flaps generated by twinPE (**Fig. 1a**). Co-transfection of plasmids encoding PE6c and twin-epegRNAs (A and B with 35-nt flaps) targeting the *AAVS1* locus, along with a linear dsDNA donor, into HEK293T cells resulted in efficient insertion of the 0.8-kb DNA (**Fig. 1b**). No insertion bands were detected in control samples, including no donor, circular plasmid template, no RT, and non-cognate pegRNA controls (**Fig. 1b**). Sequence analysis indicated that the PA approach generates accurate junctions on each side of the insertion (**Fig. 1c and Extended Data Fig. 1a)**. Analyzing the effect of flap length on PA efficiency, we observed robust editing with 30- and 50-nt flaps and very weak editing with a 13-nt flap **(Extended Data Fig. 1b)**. Using droplet digital PCR (ddPCR), we estimated the efficiency of insertion of the 0.8-kb donor into *AAVS1* to be ∼50% by PA (**Fig. 1d**). Importantly, PA was also effective for insertion of 2.2- and 4-kb donors into the *AAVS1* site in HEK293T cells (**Fig. 1e**), and of a 0.8-kb donor into the *TRAC* locus in Jurkat cells (**Extended Data Fig. 1c**).

**Fig. 1.**
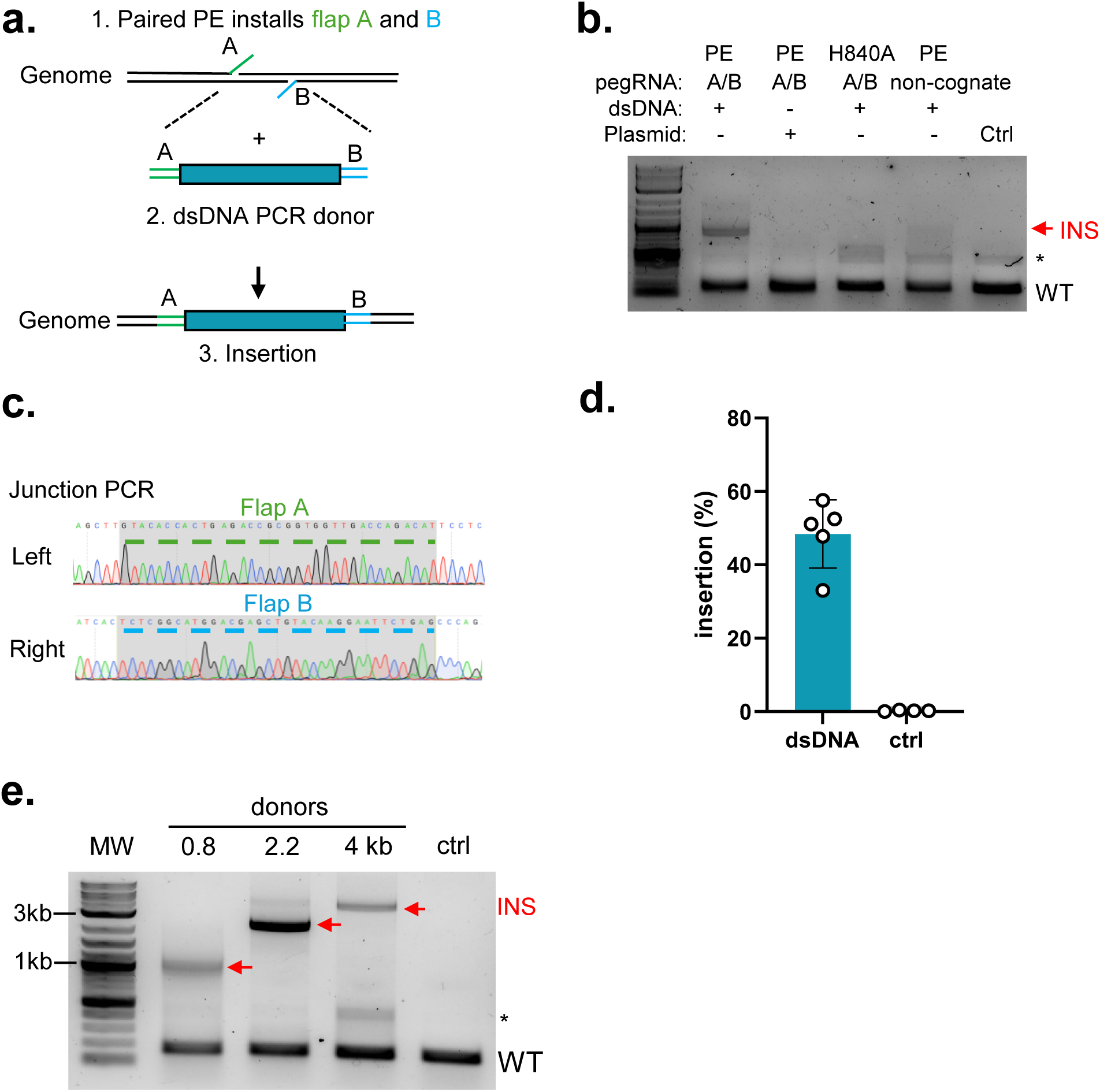
Prime assembly (PA) for large insertion using dsDNA donor. **(a)** Schematic depicting DNA insertion based on complementarity between the twinPE flaps and dsDNA donor. 35-nt flaps A and B match the ends of the dsDNA donor generated by PCR. **(b)** Agarose gel analysis of a 0.8- kb DNA donor inserted at the *AAVS1* site. HEK293T cells were transfected with the indicated components. H840A: Cas9 nickase. Plasmid: circular plasmid donor. Red arrow denotes targeted insertion (INS). Ctrl, cells without transfection. *, nonspecific band. **(c)** Sanger sequencing trace of junction regions between the donor and genome. (**d**) Insertion efficiency (% editing) of 0.8-kb donor measured by ddPCR (n = 4 to 5). **(e)** Agarose gel analysis of PA at the *AAVS1* site with donors of indicated lengths. Data and error bars indicate the mean and s.d. of independent biological replicates.

To analyze the precision of editing by PA, we performed amplicon sequencing across each insertion junction — *i.e.*, corresponding to the flap homology region. Precise junction sequences comprised 83%–88% of reads, whereas indels/SNPs comprised 12%–17% of reads (**Extended Data Fig. 2a** and **2b)**. Given the large quantification window used in this analysis (spanning the entire amplicon), some of the detected indels may result from background noise such as sequencing or PCR errors, potentially leading to an overestimation of true indel rates. To control for amplification or sequencing errors, we performed amplicon sequencing of synthetic gBlocks Gene Fragments corresponding to each junction sequence. We observed that 91%–92% of reads possessed the precise junction sequence (**Extended Data Fig. 2a** and **2b)**. Because the error rate of gBlocks Gene Fragment synthesis is estimated at 1:5000, or 0.02% (Integrated DNA Technologies), these results suggest that amplicon sequencing errors account for 8%–9% of imprecise junction sequence reads and that the editing precision might be closer to 95%. Many of the indels were consistent with MMEJ-induced deletions around the PE flap region^20,29^(**Extended Data Fig. 2e**). Nanopore long-read sequencing across the entire insertion produced similar results (**Extended Data Fig. 4a** and **4e**). Therefore, our results indicate that PA enables precise genomic insertion of large DNA fragments.

Previous studies have shown that cellular determinants and DNA repair machinery activity modulate prime editing outcomes^15,34,35^. We therefore used small molecular inhibitors to block RAD51 (B02), the Mre11-Rad50-Nbs1 (MRN) complex (Mirin), DNA-PK and PI3K (NU7026), mitosis (Nocodazole), PARP (Olaparib), Polθ (ART558), and DNA-PK (AZD-7648) to analyze how they affect PA. Among these inhibitors, the DNA-PK inhibitor AZD-7648 increased the insertion efficiency by ∼2-fold (**Fig. 2a, b** and **Extended Data Fig. 3**). PA was not affected by nocodazole, which indicates that PA is independent of cell cycle progression (**Extended Data Fig. 3**). The latter observation, along with the efficiency of PA during RAD51 inhibition, suggests that PA does not rely on the canonical HDR machinery. In cells treated with AZD-7648, PA-mediated indels decreased from 12%–17% to 9%–12%, indicating that inhibition of DNA-PK improves the accuracy of PA insertion (**Extended Data Fig. 2a** and **2b)**. To test PA at an additional genomic site, we successfully achieved GFP knock-in at exon 2 of *ACTB* and confirmed GFP expression by fluorescence microscopy (**Extended Data Fig. 5a)**. PA also replaced a 1kb *HEK3* region with a 1.3kb Halo-tag using PECas9 or PEmax (**Extended Data Fig. 5b**)^5,34^.

**Fig. 2.**
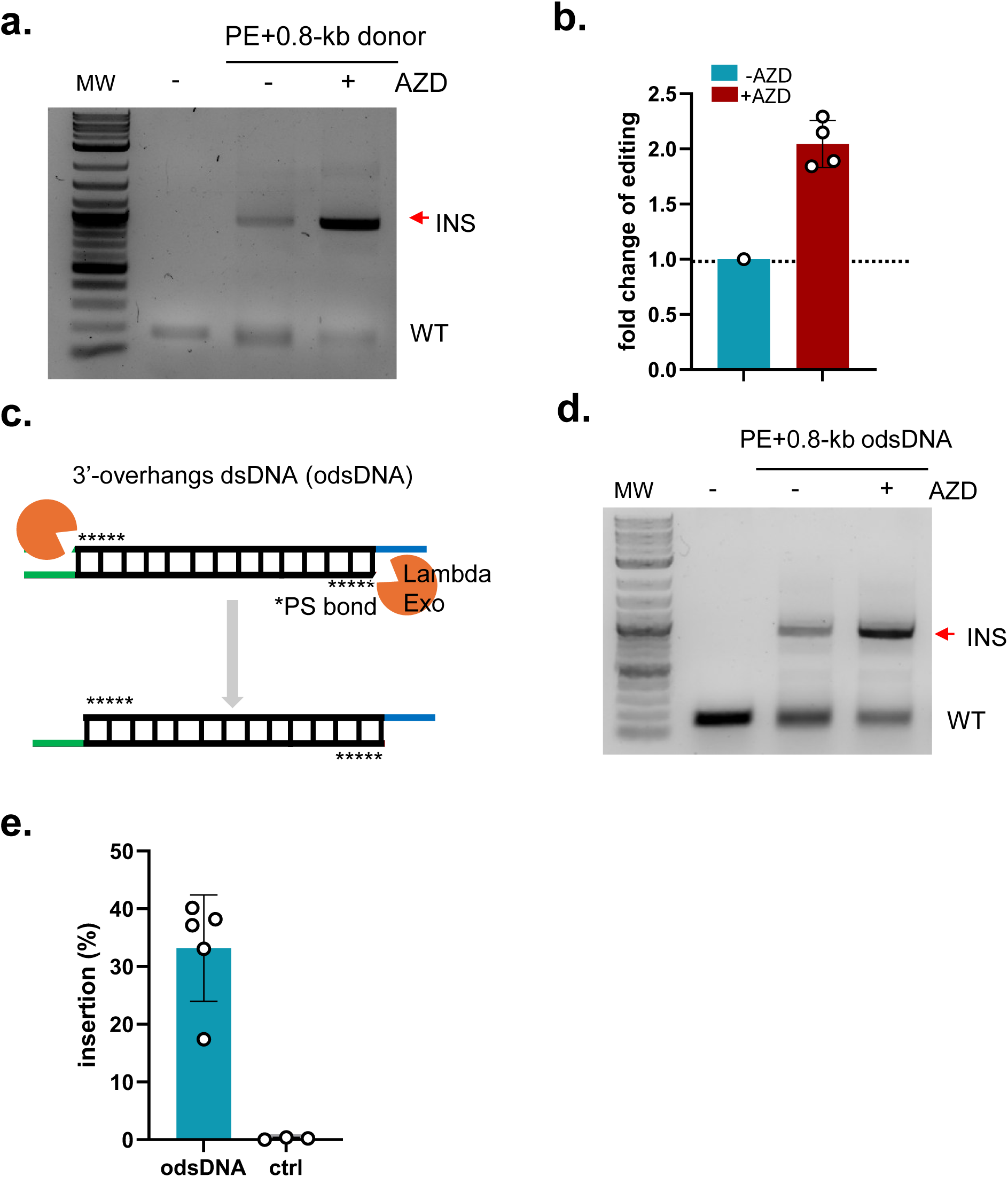
NHEJ Inhibitor (AZD-7648) and 3’-overhang dsDNA for prime assembly. **(a)** Agarose gel analysis of the 0.8-kb dsDNA insertion at the *AAVS1* site ± DNA-PK inhibitor AZD-7648. **(b)** Fold change of 0.8-kb donor insertion in the presence of AZD-7648 compared to the absence of inhibitor, measured by ddPCR (n=4).**(c)** Diagram depicting the 3′-overhang dsDNA (odsDNA) donor, generated by PCR amplification and 5′-end resection using lambda exonuclease. The 3′- overhangs anneal to the flaps generated by PE. **(d)** Agarose gel analysis of the 0.8-kb odsDNA donor insertion at the *AAVS1* site ± AZD-7648. **(e)** Insertion efficiency of 0.8-kb donor measured by ddPCR (n = 3 to 5). Ctrl = cells without transfection and treatment. Data and error bars indicate the mean and s.d. of independent biological replicates.

For the PE flaps to anneal to the DNA donor, the 5’ ends of the dsDNA donor must be displaced or resected. We therefore hypothesized that removing the 5’ ends of the 0.8-kb donor would result in a 3’ overhang dsDNA (odsDNA) donor that supports PA. We resected the 5’ ends of the 0.8-kb donor using lambda 5’-to-3’ exonuclease, generating a 3’-odsDNA donor (**Fig. 2c**). Five internal phosphorothioate (PS) linkages adjacent to the flap homology region were introduced to prevent excessive 5’-end resection (**Fig. 2c**). The 0.8-kb odsDNA supported 20%–40% insertion at *AAVS1* by PA (**Fig. 2d** and **2e)**. Compared to the unmodified dsDNA donor, the odsDNA donor slightly reduced the frequency of insertion (**Figs. 1d** and **2e)** but resulted in slightly increased precision (**Extended Data Fig. 4e**). Notably, treatment with DNA-PK inhibitor AZD-7648 substantially reduced the frequency of indels in odsDNA-mediated insertions to the background indel levels from the gBlock control (**Extended Data Fig. 2c, 2d**, and **4e**), which is consistent with the results of dsDNA with inhibitor treatment. We further showed that PA mediates a 95 bp insertion containing an I-SceI site using two single-stranded DNA (ssDNA) donors (**Extended Data Fig. 6)**.

To overcome the donor synthesis limitations imposed by long-range PCR, we tested the feasibility of combinatorial donor assembly by dividing the 0.8-kb dsDNA donor into two segments that overlap by 30-bp, mimicking Gibson assembly^36^ (**Fig. 3a**). The split donor supported the PA-mediated 0.8-kb insertion with ∼30% efficiency without AZD-7648 (**Fig. 3a-c**). Additionally, AZD- 7648 treatment increased the insertion efficiency and accuracy (**Fig. 3b,d**). In the absence of AZD-7648, we observed 6.5% indels, most of which corresponded to NHEJ events between the two donor segments. However, indels were reduced to 2% in the presence of AZD-7648 (**Fig. 3d**), consistent with the role of DNA-PK in NHEJ^37,38^. Sanger sequencing confirmed that the donor segments were correctly assembled and inserted in cells (**Fig. 3e**).

**Fig. 3.**
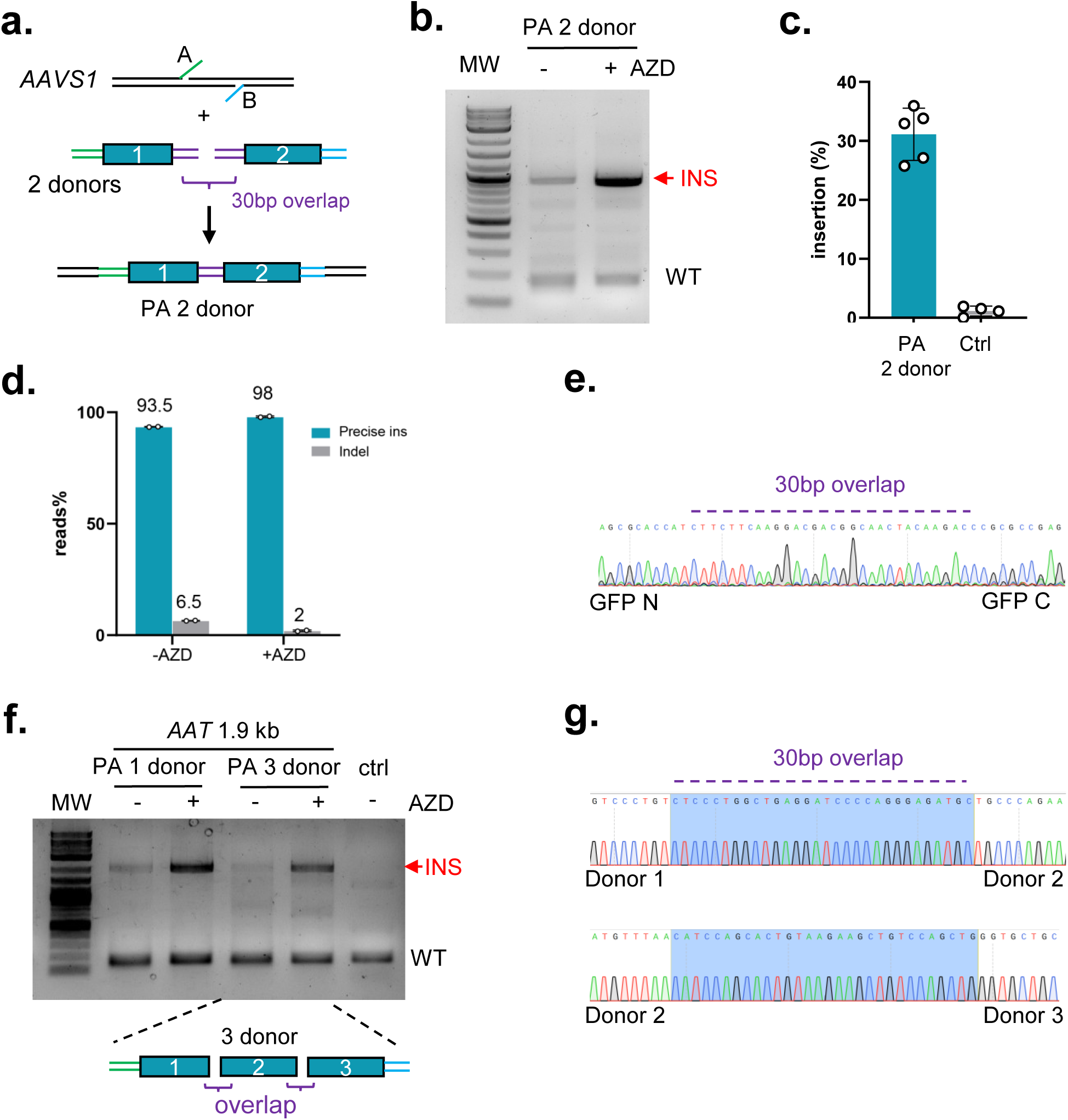
Multiple donor prime assembly in mammalian cells. **(a)** Illustration of PA insertion of two donor segments that overlap by 30-bp (purple). The outside ends of dsDNA donor (green and blue) match the two 35-nt flaps generated by twinPE. **(b)** Agarose gel analysis of insertion mediated by PA 2 donor ± AZD-7648. **(c)** Insertion efficiency of PA 2 donor or control measured by ddPCR (-AZD). HEK293T cells were transfected with PE6, pegA+pegB, GFP-N (432 bp) and GFP-C (431 bp) PCR donors (n = 4 to 5). Ctrl = cells without transfection and treatment. **(d)** Average percent reads (n = 2 biological replicates) that correspond to precise insertion and indels (i.e., NHEJ insertion) between the GFP-N and GFP-C donors ± AZD-7648. **(e)** Sanger sequencing trace of the junction regions between the GFP-N and GFP-C donor segments. **(f)** PA 3 donor of a 1.9-kb IRES-*AAT* insertion at the *AAVS1* site. 293T cells were transfected with PA 1 donor or 3 donor ± AZD-7648. **(g)** Nanopore sequencing confirms accurate assembly of PA 3 donors (+AZD). 30bp overlap sequences are highlighted. Data and error bars indicate the mean and s.d. of independent biological replicates.

We next explored the assembly of three overlapping segments of a 1.9-kb alpha-1 antitrypsin (*AAT*) gene into the *AAVS1* locus. Successful integration was observed using three-segment PA (**Fig. 3f**). Consistent with results from two-segment PA, insertion efficiency improved upon AZD- 7648 treatment (**Fig. 3f**). Nanopore sequencing confirmed that insertion is precise (**Fig. 3g**). These results show that our approach effectively mimics Gibson assembly within mammalian cells^36^, advancing the ability to precisely insert large DNA segments into the mammalian genome.

Next, we tested whether PA could mediate the insertion of a full-length *Duchenne Muscular Dystrophy* (*DMD*) gene (11.3 kb) into the *AAVS1* locus in HEK293T cells (**Fig. 4a)**. We co-transfected a column-purified, 11.3-kb *DMD* PCR donor (**Fig. 4b**), PE6c, and twin-epegRNAs into HEK293T cells. After three days we harvested cells and isolated genomic DNA (gDNA), amplified the insertion junctions, and confirmed accurate insertion of the *DMD* donor by sequencing (**Fig. 4c-d**). Agarose gel analysis using outward-facing primers, which flank the insertion site to amplify the entire insertion, further validated the full-length 11.3-kb insertion (**Fig. 4e**). Insertion was enhanced upon AZD-7648 treatment, consistent with previous results (**Fig. 4e**).

**Fig. 4.**
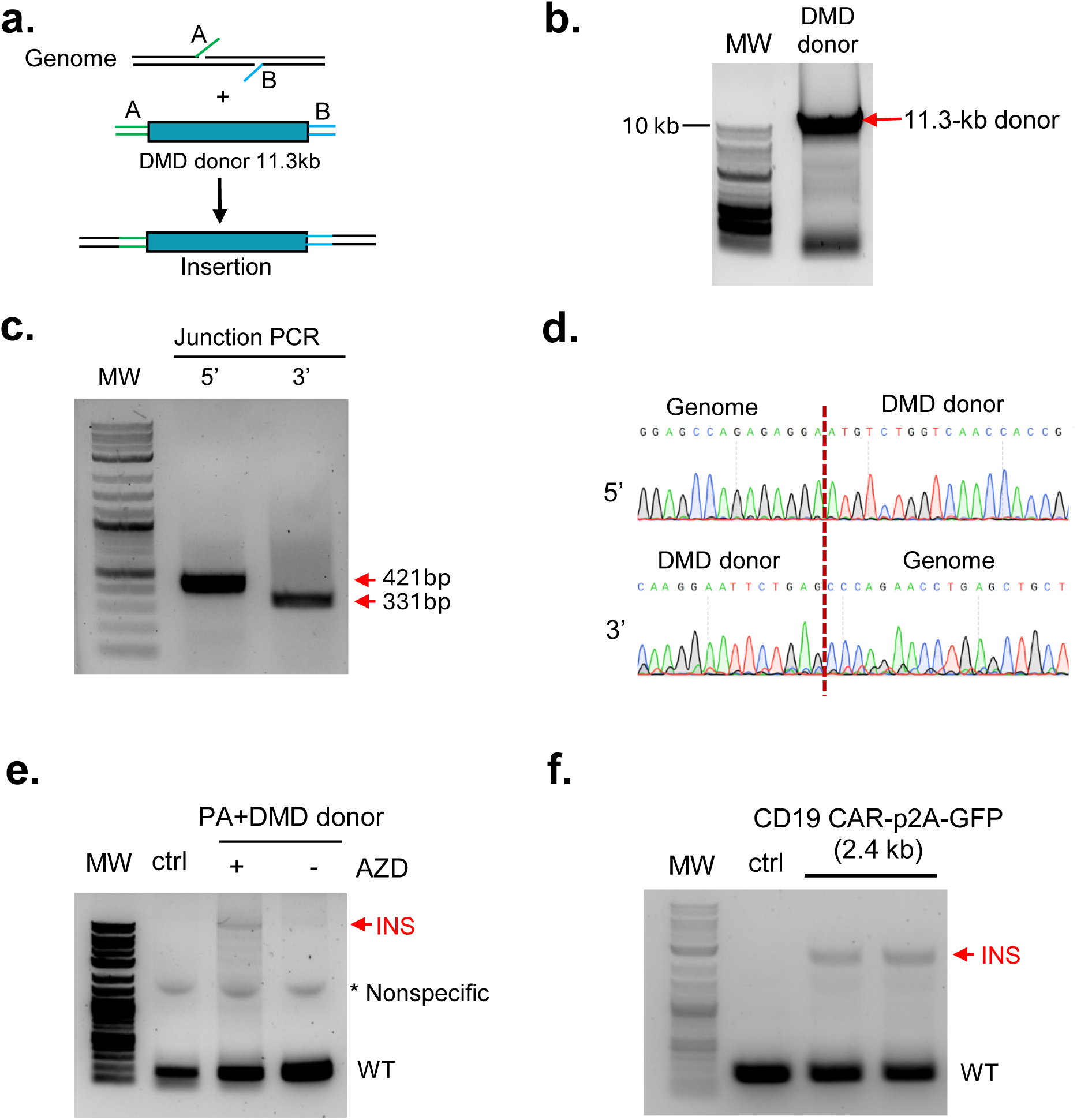
Prime assembly for *DMD* and CAR insertion. **(a)** Schematic of PA insertion of the 11.3-kb full length *DMD*. **(b)** DMD donor was generated by PCR. **(c)** Agarose gel of genome– *DMD* insertion junction PCR. Expected band sizes are indicated. **(d)** Sanger sequencing traces of the junction PCR between the genome and donor. **(e)** Agarose gel analysis of 11.3-kb *DMD* insertion at the *AAVS1* site in the absence or presence of DNA-PK inhibitor AZD-7648. HEK293T cells were transfected with PE6c, pegA+pegB, and an 11.3-kb *DMD* donor. Agarose gel of the *DMD* gene insertion at the *AAVS1* site. Ctrl = cells without transfection and treatment. **(f)** Agarose gel of CD19 CAR-p2A-GFP insertion into the T-cell receptor α constant (*TRAC*) locus in 293T cells in the presence of AZD-7648. Replicate transfections are shown.

Lastly, we tested whether a CD19-specific CAR gene could be efficiently inserted into the T-cell receptor α constant (*TRAC*) locus in HEK293T cells (**Fig. 4f**)^39^. Our results confirmed the successful PA integration of the 2.4 kb CD19 CAR-P2A-GFP sequence into the *TRAC* locus, demonstrating that PA enables the integration of large DNA cargo at therapeutically relevant sites.

## Discussion

In this study, we demonstrated that PA, using twinPE and linear dsDNA or ssDNA donors, enables large insertions ranging from 0.1 to >11 kb in mammalian cells. The efficiency was up to 50% for a 0.8kb donor. We also found that 3’-overhang dsDNA donors support PA-mediated large insertions. Furthermore, our combinatorial donor assembly design overcomes the challenge of producing long DNA donors and enables the simultaneous and combinatorial assembly of multiple donors within mammalian cells. The PA system is cost-effective, scalable, and adaptable, making it a powerful tool for large gene insertions, replacements, and precise genome modifications in mammalian cells.

Although the integrase/recombinase-independent nature of our PA system offers potential delivery advantages over previous recombinase-based strategies, future studies will be needed to explore effective non-viral *in vivo* delivery methods for PA (similar to most other targeted long- insertion technologies). Given that various chemical modifications can be introduced into linear DNA donors—e.g., via PCR primers—further optimization could increase insertion efficiency while minimizing toxicity and immunogenicity of dsDNA donors for *in vivo* delivery^40^. Additionally, an affinity module that tethers donor DNA to the editing site might further improve PA efficiency^20,28^.

The precise DNA synthesis and repair mechanisms underlying PA remain to be defined. We consider it likely that PA insertions are driven by endogenous DNA polymerases, but the identities of any such polymerases will require additional studies. Results with overlapping split donors strongly imply that the dsDNA serves as a template, not a direct ligation substrate. Given the involvement of single-stranded PE flaps and linear dsDNA, it is reasonable to hypothesize that PA-related repair pathways differ from traditional HDR, NHEJ, and PE-associated mechanisms^24,34,41,42^. Our findings with DNA repair inhibitors support this speculation, but further studies are needed to elucidate the repair pathways involved^34^. Although AZD-7468 can enhance PA efficiency, this benefit comes with trade-offs based on undesirable effects of this inhibitor that have been observed^43^.

Recent advances in PE have opened new possibilities for precise large insertions, but the efficiency of inserting >400-bp donors remains low^5,6,29,30,32,44^. By leveraging programmable flaps generated by PE, our PA system enables controlled and precise insertions without relying on double-stranded breaks (DSBs), long homology arms, or additional recombinase elements. This provides a potentially versatile approach for genome editing and gene therapy.

## Materials and Methods

### Cell culture and small molecule inhibitor treatment

HEK293T cells and Jurkat cells were acquired from ATCC and cultured at 37 °C with 5% CO2 in Dulbecco’s modified Eagle medium (DMEM) with 10% fetal bovine serum (FBS) and 1% penicillin/streptomycin. Six hours post-transfection, HEK293T cells were treated using inhibitors with indicated concentrations. The Inhibitors B02, Mirin, NU7026, AZD-7648, and Nocodazole were purchased from MedChemExpress, and Olaparib was purchased from Selleck Chemicals. ART558 was purchased from TargetMol.

### Plasmids

epegRNA expression plasmids were constructed using a custom vector (BfuAI- and EcoRI-digested) that has been described elsewhere^45^. The gBlocks gene fragments with homology sequence to the vector were synthesized by Integrated DNA Technologies (IDT), followed by Gibson assembly using Gibson Assembly Master Mix (New England Biolabs). The donor plasmids were constructed by Gibson assembly using pMD217 Vector. Colonies were selected and further confirmed by Sanger using the commercial human U6, M13R and other specific primers. Sequences of epegRNA are listed in **Supplementary Table 1**.

The SA-GFP donor plasmid used for double-stranded DNA (dsDNA) donor preparation was constructed using the pMD217 vector. The human U6 promoter was removed, and the gene sequence of interest was inserted via Gibson assembly.

pCMV-PE6c was a gift from David Liu (Addgene plasmid #207853). p37-2iDMD-LR was a gift from Michele Calos (Addgene plasmid #88892). pSLCAR-CD19-28z was a gift from Scott McComb (Addgene plasmid #135991).

### Linear dsDNA/ssDNA Donor Preparation and transfection

For linear dsDNA purification, Phusion Flash High-Fidelity PCR Master Mix (Thermo Scientific™) was used for PCR amplification according to the manufacturer’s instructions. To obtain a high concentration for transfection, a total of 0.5 mL of PCR reaction product was purified using two QIAquick PCR Purification (QIAGEN) columns with the same elution buffer. DNA was eluted in 50 µL of nuclease-free water. Specific primer sequences for detecting insertions and preparing donors are listed in **Supplementary Table 2.**

For odsDNA preparation, PCR products were digested with lambda exonuclease (New England Biolabs, NEB) at 37 °C for 3 hours, followed by heat inactivation at 80 °C, as per the manufacturer’s protocol. The digested products were subsequently purified using QIAquick PCR Purification columns. Donor sizes were assessed via agarose gel electrophoresis, and DNA concentration was measured using a NanoDrop™ One UV-Vis Spectrophotometer.

The ssDNA syntheses were performed by Genewiz from Azenta Life Sciences. dsDNA/ssDNA donor sequences and plasmid template sequences were listed in **Supplementary sequences**.

### Transfection and Genomic DNA Isolation

Cell transfection was performed using the Lipofectamine 3000 reagent (Invitrogen, L3000015) according to the manufacturer’s instructions. Briefly, 0.8 × 10⁵ cells were seeded per well in a 12- well plate and incubated overnight. Transfection was carried out using 3.5 μL of Lipofectamine 3000 and P3000 reagent (2 μL per μg of DNA). For each well, 500 ng of epegRNA plasmids, 1 μg of PE6 plasmids, and 1–2 μg of purified linear DNA donors or 100 pmol ssDNA were used.

Seventy-two hours post-transfection, cells were collected and lysed using 100 μL of Quick Extraction Buffer (Lucigen). The lysate was incubated in a thermocycler at 65 °C for 15 minutes, followed by 98 °C for 5 minutes, according to the Quick Extraction protocol.

### Nucleofection

For Jurkat cells nucleofection, the Neon electroporation system was used. Briefly, 1× 10⁵ cells were used for each electroporation with 10-μl Neon tips. For each electroporation, 500 ng of epegRNA plasmids, 1 μg of PE6 plasmids, and 1 μg of purified linear DNA donors were used with the following electroporation parameters: 1,600 V, 10 ms, three pulses. After electroporation, cells were plated in prewarmed 48-well plates with DMEM containing 10% FBS and inhibitors and incubated for 72 h before analysis. Genomic DNA isolation was performed using 50 μL of Quick Extraction Buffer (Lucigen) according to the Quick Extraction protocol.

### Sanger sequencing

PCR amplification was performed at the target locus using Phusion Flash PCR Master Mix (Thermo Fisher) and specific primers. The PCR products were purified using agarose gel electrophoresis, and Sanger sequencing was carried out by Genewiz (Azenta Life Sciences).

### Quantification of DNA insertion efficiency via droplet digital PCR (ddPCR)

A ddPCR approach was used to quantify the efficiency of precision insertion junction in comparison to a reference amplicon. Briefly, genomic DNA was mixed with the ddPCR Supermix (no dUTP; Bio-Rad), along with probes (250 nM) and primers (900 nM), in a final volume of 20 μl. Droplet generation was performed using a QX200 Manual Droplet Generator (Bio-Rad). The following PCR program was used: 95 °C for 10 min, 35 cycles of 94 °C for 30 s and 58 °C for 1 min, 98 °C for 10 min, and 4 °C holds. Data acquisition was performed on a QX200 Droplet Reader (Bio-Rad). Data analysis was performed using QuantaSoft (Bio-Rad). Editing efficiency was calculated by using the following formula: editing % = 100% × FAM+/ HEX+. Sequences of probes and primers are listed in **Supplementary Table 4.**

### Amplicon sequencing and data analysis

Sequencing library preparation has been described previously^45^. Briefly, for the first round of PCR, the specific primers with Illumina forward and reverse adapters (**Supplementary Table 3**) were used to amplify the genomic sites of interest by Phusion Hot Start II PCR Master Mix (Thermo Scientific). For the second round of PCR, primers containing unique Illumina barcodes were used and PCR reactions were performed using the following parameters: 98 °C for 10 s, 20 cycles of 98 °C for 1 s, 50 °C for 5 s and 72 °C for 7 s, followed by 72 °C for 2 min as a final extension. The DNA products of second-round PCR were collected and purified by gel purification using the QIAquick Gel Extraction kit (Qiagen), and DNA concentration was determined by Qubit dsDNA HS assay. Subsequently, the library was sequenced on an Illumina MiniSeq following the manufacturer’s protocol.

To quantify precise edits and indels at insertion junctions, CRISPResso2 was run in NHEJ mode with expected junction sequences after editing using default settings except “ignore_substitutions” set to “TRUE” and “qwc” (quantification window) set to include the full amplicon except the first and last 15 nucleotides. Percentage of “UNMODIFIED” reads reported by CRISPResso2 is designated as the frequency of perfect editing and percentage of “MODIFIED” reads is designated as the frequency of indel.

### Nanopore sequencing and data analysis

PCR products were purified using 0.7 x homemade magnetic beads to eliminate the short fragment amplified from the unedited genome^46^. The purified DNA was quantified by NanoDrop and sent to Plasmidsaurus for their “Premium PCR” service (ligation-based long-read nanopore sequencing). Raw reads in FASTQ format were requested and used for subsequent analysis.

Raw reads were aligned to the expected sequences after editing using minimap2 with the parameter “-x map-ont”^47,48^. The output BAM files were filtered for reads with >10% homology compared to the reference. Locations and sizes of the indels were extracted from the filtered BAM files. “Flap indel” frequency was calculated using (number of reads with indels >6 bp in the 100 bp region flanking the insertion point) / (total number of reads after filtering). “Precise editing” frequency was calculated using 1 – (flap indel).

## Data availability

High-throughput sequencing data are being deposited to the NCBI Sequence Read Archive database.

### Author contributions

B.L., E.J.S., and W.X. conceptualized the project. B.L., A.P., X.Z., T.J., and L.Z. conducted molecular biological and cellular experiments. B.L. and H.C. conducted high-throughput sequencing and bioinformatic analyses. B.L., E.J.S., and W.X. interpreted the data and wrote the paper with all co-authors.

## Supporting information

Supplementary Tables, sequences, and notes.

## Acknowledgments

We thank the Xue and Sontheimer labs for helpful discussions. We thank D. Conte for editing the manuscript and S. DeLeon for excellent contributions to this project. We also thank Sébastien Levesque and Dan Bauer for communication of unpublished results. This work was supported by grants from the National Institutes of Health (R01CA275945, R01GM150279, P01HL158506 and UH3HL147367), and the Leducq Foundation Transatlantic Network of Excellence Program. T.J. was supported by grants from the National Institutes of Health (R00HL153940) and Cystic Fibrosis Foundation (005363I223).

---

**Extended Data Fig 1.**
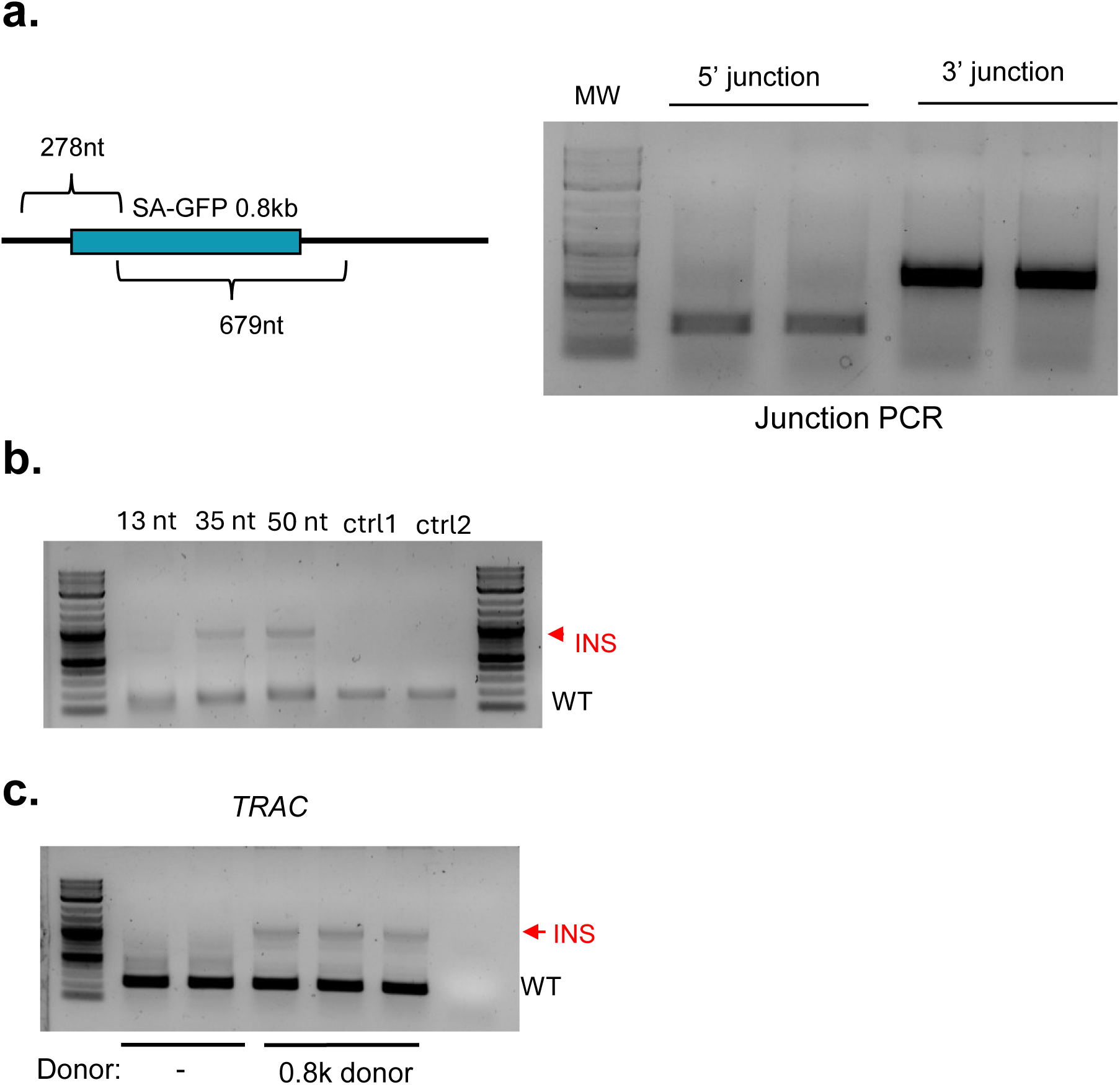
Junction PCR and different flap lengths for PA. **(a)** Junction PCR and agarose gel analysis of PA insertion. Replicate transfections are shown. **(b)** A comparison of varying flap lengths for PA insertion. HEK293T cells were transfected with PE6, pegA+pegB, and dsDNA donors. ctrl1 = cells transfected with GFP plasmid; ctrl2 = cells without transfection and treatment. **(c)** 0.8k insertion at the *TRAC* locus in Jurkat cells.

**Extended Data Fig 2.**
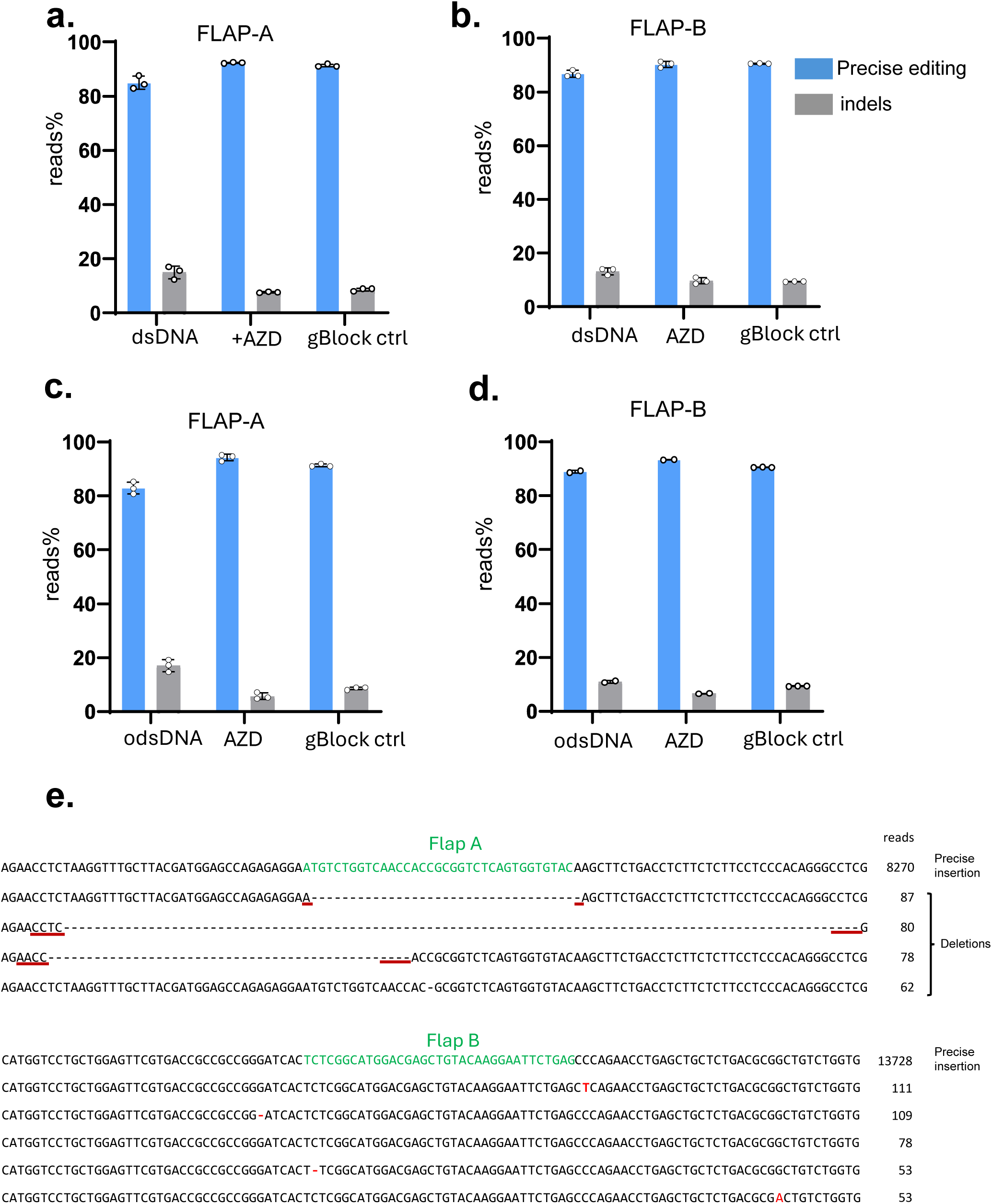
Evaluating the accuracy of PA insertions by amplicon sequencing. **(a-b)** Amplicon sequencing analysis of two flap-junction regions using dsDNA donor ± AZD inhibitor (n=3). **(c-d)** Amplicon sequencing analysis of two flap-junction regions using odsDNA donor ± AZD inhibitor (n = 2 to 3). **(e)** Representative amplicon sequencing reads for flap-junction regions. Some deletions near flap A are likely caused by short homology (underlined). The top five reads are shown. Data and error bars indicate the mean and s.d. of independent biological replicates.

**Extended Data Fig 3.**
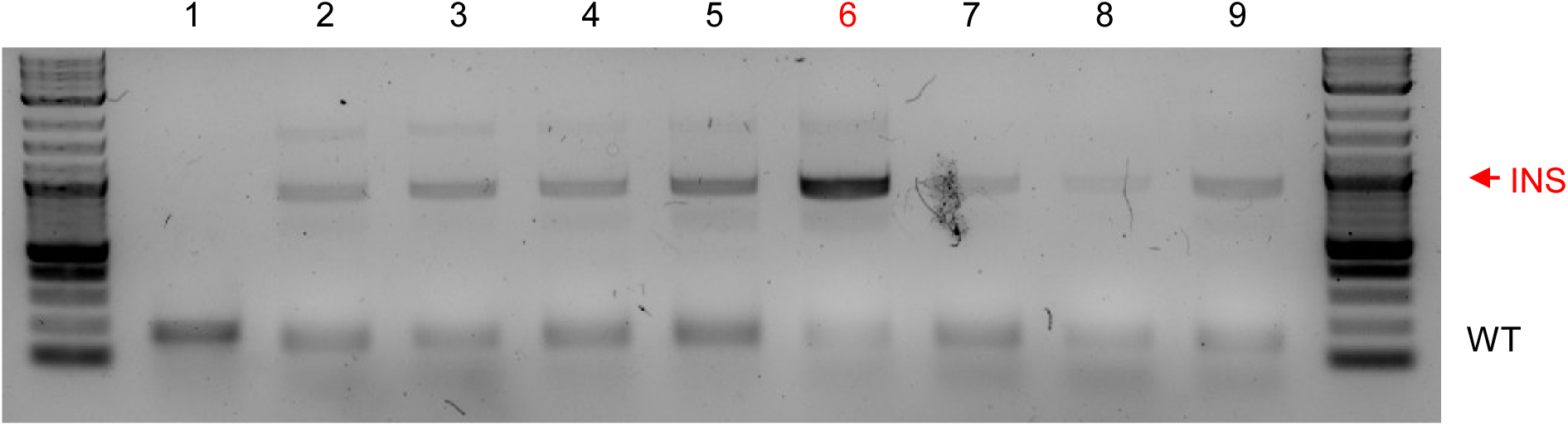
PA insertion (*AAVS1* site, 0.8kb donor) upon inhibitor treatment. 1. No donor control, 2. PA with no drug treatment, 3. RAD51 Inhibitor B02 (10 μM), 4. Mre11- Rad50-Nbs1 (MRN) complex inhibitor Mirin (10 μM), 5. DNA-PK and PI3K inhibitor NU7026 (1 μM), 6. DNA-PK inhibitor AZD-7648 (1 μM), 7. M phase inhibitor Nocodazole (200 nM), 8. PARP inhibitor Olaparib (5 μM), 9. PolΘ inhibitor ART558 (200 nM).

**Extended Data Fig 4.**
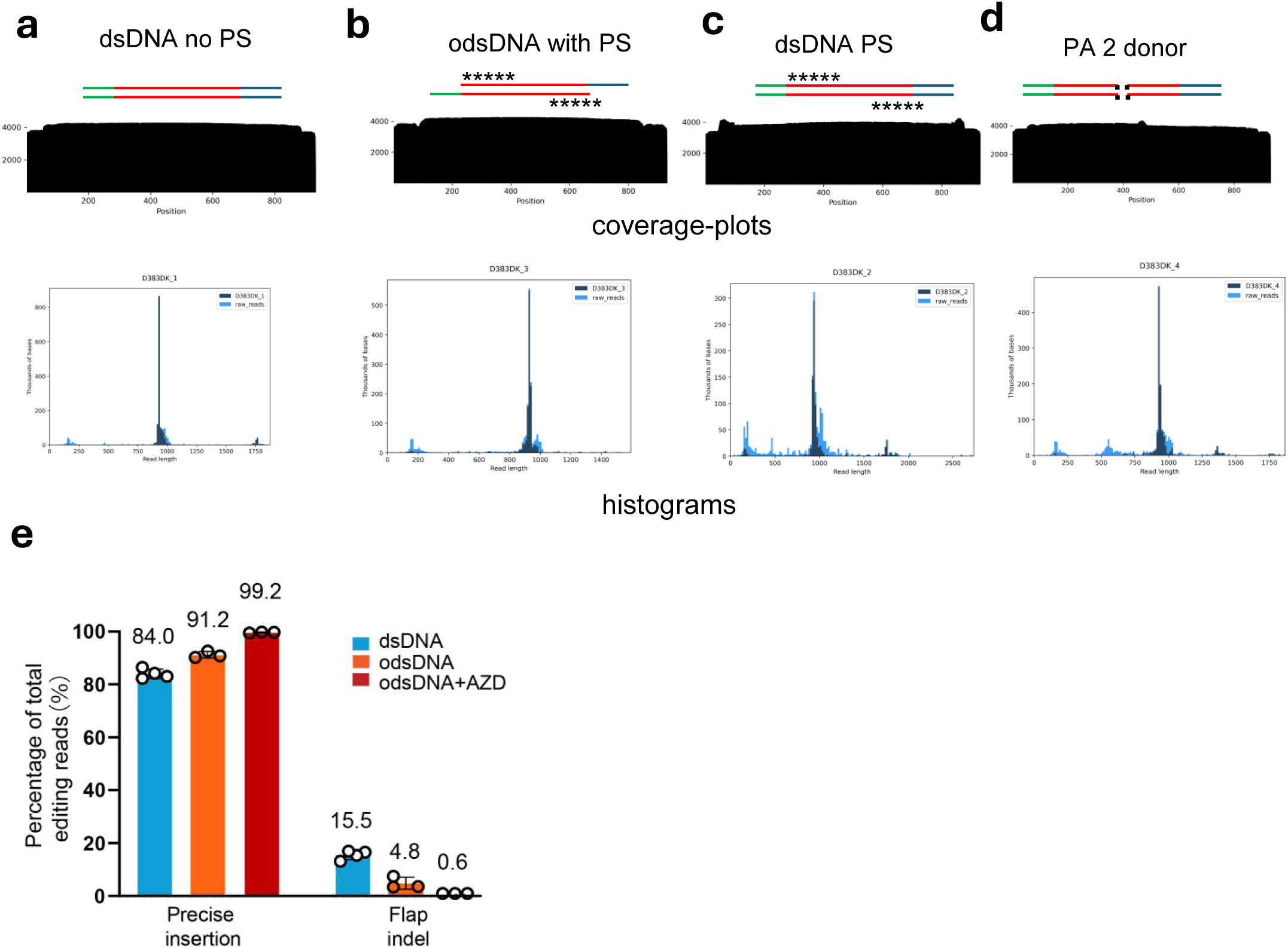
Evaluation of the accuracy of PA insertions. **(a-d)** Representative coverage plots and histograms from nanopore sequencing. The coverage plots display the relative sequencing coverage at each position of the consensus sequence. The histogram displays the lengths of the raw reads produced, with read length (bp) on the x-axis and thousands of bases of data collected (kb) at that length on the y-axis. **(e)** Nanopore sequencing analysis of PA (n=3 to 4). Data and error bars indicate the mean and s.d. of independent biological replicates.

**Extended Data Fig 5.**
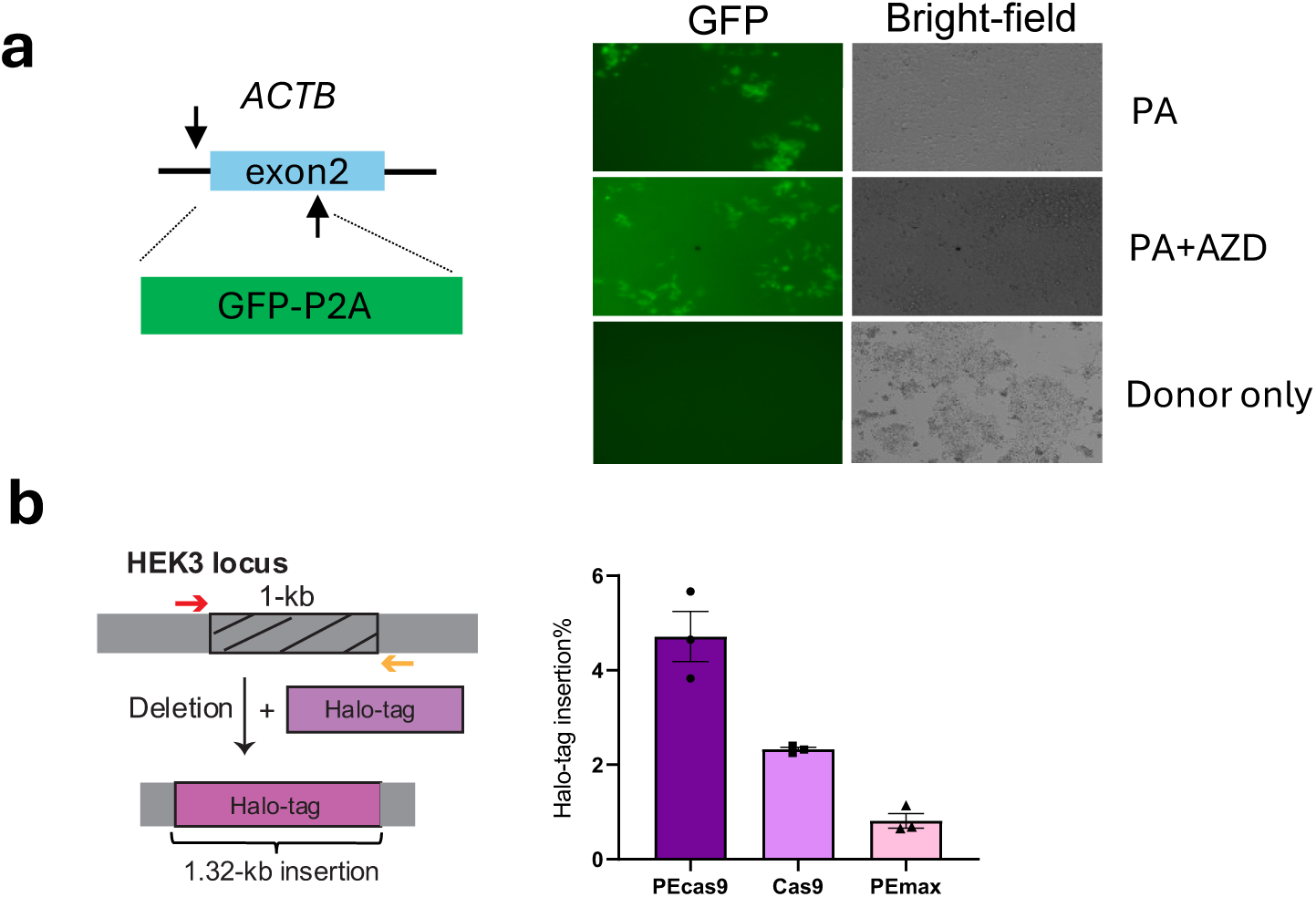
PA insertion at *ACTB* and HEK3 sites. **(a)** Left, Schematic of GFP insertion into *ACTB* by PA. Right, Fluorescence and bright-field images showing functional GFP insertion into the *ACTB* site 5 days after transfection. **(b)** Replacement of a 1-kb target sequence with a HaloTag at the *HEK3* site by PA using PECas9 or PEmax (n = 3). Data and error bars indicate the mean and s.d. of independent biological replicates.

**Extended Data Fig 6.**
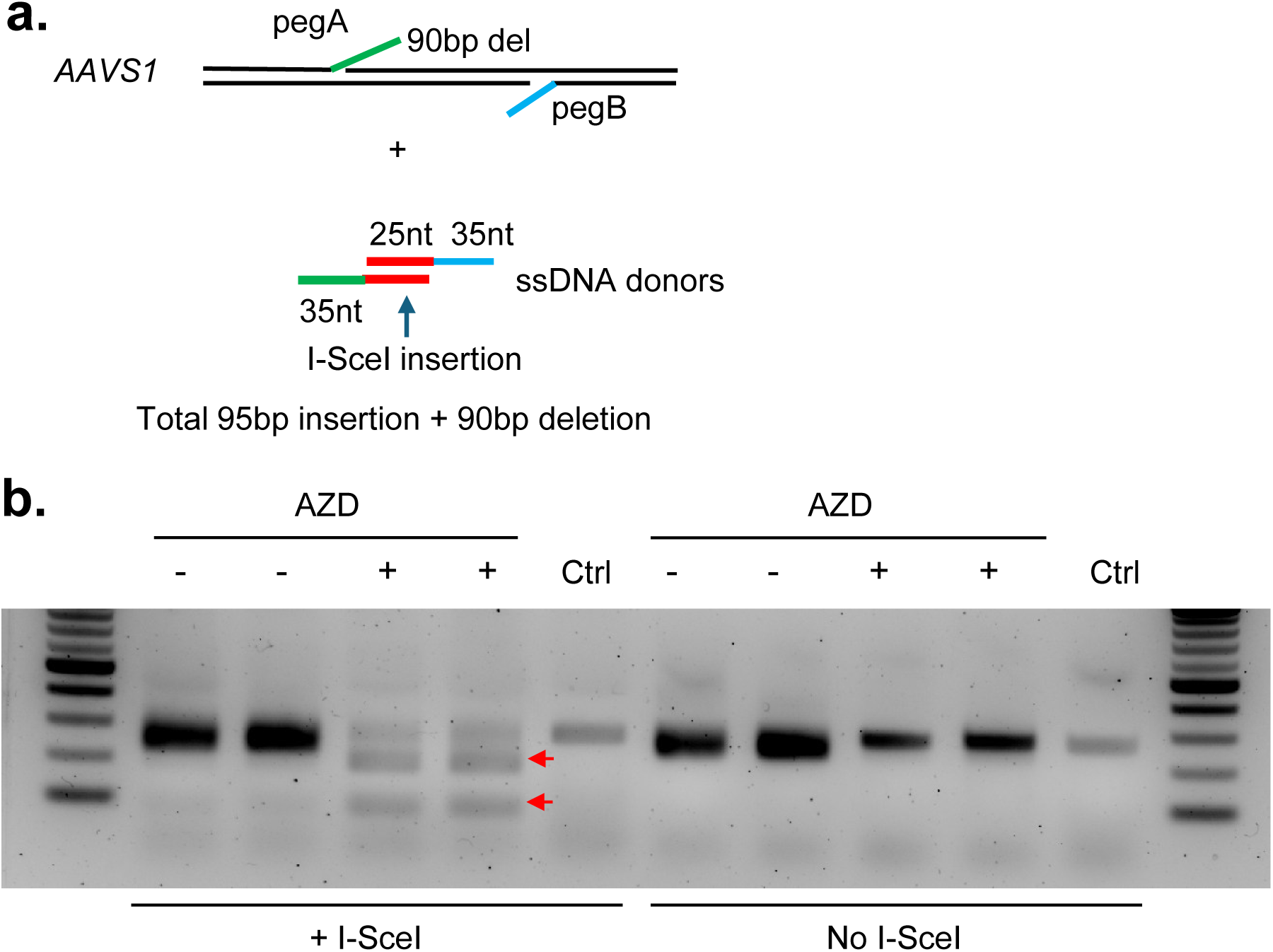
PA mediates a 95 bp insertion containing an I-SceI site using single-stranded DNA (ssDNA) donors. **(a)** Schematic of PA with ssDNA donors. The 25nt insertion contains I-SceI site. **(b)** HEK293T cells were transfected with PE6c, pegRNAs and two ssDNA donors. Genomic PCR products were treated with I-SceI. The cleavage bands indicate insertion (red arrows). AZD increased insertion efficiency. Ctrl: cells without transfection.

