## Supplementary Tables, sequences, and notes. for "Prime assembly with linear DNA donors enables large genomic insertions"

**Supplementary Information**

**Supplementary Table 1.** Twin epegRNAs.

**Supplementary Table 2**. Primers used for PCR.

**Supplementary Table 3**. Primers used for high throughput sequencing.

**Supplementary Table 4**. Primers used for droplet digital PCR (ddPCR).

**Supplementary Notes**

**Supplementary sequences**. dsDNA/ssDNA donor sequences and plasmid template sequences.

**Supplementary Table 1**. Twin epegRNAs.

| epegRNA | spacer | RTT | PBS |
|---|---|---|---|
| *AAVS1_peg1* | GATGGAGCCAGAGAGGATCC | ggccctgtgggaggaagagaagaggtcagaagctt | TCCTCTCTGGCTC |
| *AAVS1_peg1B* | GATGGAGCCAGAGAGGATCC | GTACACCACTGAGACCGCGGTGGTTGACCAGACAT | TCCTCTCTGGCTC |
| *AAVS1_peg2* | GCAGCTCAGGTTCTGGGAGA | TCTCGGCATGGACGAGCTGTACAAGGAATTCTGAG | CCCAGAACCTGAG |
| *AAVS1_peg3* | GCAGCTCAGGTTCTGGGAGA | CCGCGGTGGAGCTCCAATTCGCCCTATAGTGAGTC | CCCAGAACCTGAG |
| *AAVS1-13nt_peg1* | GATGGAGCCAGAGAGGATCC | AGGTCAGAAGCTT | TCCTCTCTGGCTC |
| *AAVS1-13nt_peg2* | GCAGCTCAGGTTCTGGGAGA | AAGGAATTCTGAG | CCCAGAACCTGAG |
| *AAVS1-50nt_peg1* | GATGGAGCCAGAGAGGATCC | TGCCAGATCTCTCGAGGCCCTGTGGGAGGAAGAGAAG AGGTCAGAAGCTT | TCCTCTCTGGCTC |
| *AAVS1-50nt_peg2* | GCAGCTCAGGTTCTGGGAGA | CGCCGCCGGGATCACTCTCGGCATGGACGAGCTGTAC AAGGAATTCTGAG | CCCAGAACCTGAG |
| *TRAC_peg1* | Eyquem, J. et al. (2017) | GTACACCACTGAGACCGCGGTGGTTGACCAGACAT | TGCCCCAAC |
| *TRAC_peg2* | Eyquem, J. et al. (2017) | TCTCGGCATGGACGAGCTGTACAAGGAATTCTGAG | GATATCCAGAACC |
| *ACTB_peg1* | GCTATTCTCGCAGCTCACCA | GTGAACAGCTCCTCGCCCTTGCTCACCATGGTGGC | TGAGCTGCGAGAA |
| *ACTB_peg2* | TGGAGGGGAAGACGGCCCGG | TGCAAGGCCGGATTTGCAGGCGATGATGCTCCACG | GGCCGTCTT |
| *ACTB_peg3* | TGGAGGGGAAGACGGCCCGG | TGTAAAGCTGGTTTTGCAGGTGATGACGCTCCTCG | GGCCGTCTT |

**Supplementary Table 2**. Primers used for PCR.

**2.1** Primers of genomic PCR for insertion detection

| Gene locus | F (5'-3') | R (5'-3') |
|---|---|---|
| *AAVS1* | CTTGCCAGAACCTCTAAGGT | CCAGGATCAGTGAAACGCAC |
| *TRAC* | CACGAGCAGCTGGTTTCTAA | ATCGGTGAATAGGCAGACAGAC |
| *ACTB* | CGACCTCGGCTCACAGCG | CCACCCAGCCAGCTCCC |

**2.2** Primers for making dsDNA donors

| Gene locus | F (5'-3') | R (5'-3') |
|---|---|---|
| *AAVS1* 0.8k dsDNA | ATGTCTGGTCAACCACCGCGGTCTCAGTGGTGTA Caagcttctgacctcttctcttcc | CTCAGAATTCCTTGTACAGCTCGTCCATGCCGAGAGTG ATCCCGGCGGCGGTC |
| *AAVS1 2.2*k dsDNA | ATGTCTGGTCAACCACCGCGGTCTCAGTGGTGTA Caagcttctgacctcttctcttcc | GACTCACTATAGGGCGAATTGGAGCTCCACCGCGGCC ACTGAGCGTCAGACCC |
| *AAVS1* 4k dsDNA | ATGTCTGGTCAACCACCGCGGTCTCAGTGGTGTA Caagcttctgacctcttctcttcc | GACTCACTATAGGGCGAATTGGAGCTCCACCGCGGGG CTCTCTCTCTCTTGACGC |
| *AAVS1-0.8k-two-segments-junction* | GTCTTGTAGTTGCCGTCGTCCTTGAAGAAG | CTTCTTCAAGGACGACGGCAACTACAAGAC |
| *AAVS1-AAT-1.9k* | aagcttctgacctcttctcttcctcccacagggccTGTAAGTAGCCT GACTAGCCTAG | ctcagaattccttgtacagctcgtccatgccgagacctgcaggctacaggtcc |
| *AAVS1-AAT-1.9k-Three-segments-Junc1* | gcatctccctggggatcctcagccagggag | ctccctggctgaggatccccagggagatgc |
| *AAVS1-AAT-1.9k-Three-segments-Junc2* | cagctggacagcttcttacagtgctggatg | catccagcactgtaagaagctgtccagctg |
| *TRAC 0.87*k dsDNA | ATGTCTGGTCAACCACCGCGGTCTCAGTGGTGTA C | CTCAGAATTCCTTGTACAGCTCGTCCATGCCGAGA |
| *TRAC* 2.6k dsDNA | ATGTCTGGTCAACCACCGCGGTCTCAGTGGTGTA Cgactacaaggatcatgacggagact | GACTCACTATAGGGCGAATTGGAGCTCCACCGCGGcgc gttacttgtacagctcgtccat |
| *ACTB peg1 + peg2 dsDNA* | gccaccatggtgagcaagg | CGTGGAGCATCATCGCCTGCAAATCCGGCCTTGCAcata ggaccagggttttcctcg |
| *ACTB peg1 + peg3 dsDNA ROUND1* | gccaccatggtgagcaagg | CCAGAACCATTATCCACTACTAATGCAGCTATGTCGTC GTCcataggaccagggttttcc |
| *ACTB peg1 + peg3* dsDNA ROUND2 | gccaccatggtgagcaagg | CGAGGAGCGTCATCACCTGCAAAACCAGCTTTACACAT TCCAGAACCATTATCCACTAC |

### **2.3** Primers for making odsDNA donor

*denotes phosphorothioate (PS) linkage

**primers are 5’ phosphorylated

| Gene locus | F (5'-3') | R (5'-3') |
|---|---|---|
| *AAVS1* 0.8k dsDNA | ATGTCTGGTCAACCACCGCGGTCTCAGTGGTGTACa*a*g*c *t*tctgacctcttctcttcc | ctcagaattccttgtacagctcgtccatgccgagag*t*g*a*t* cccggcggcggtc |

### **2.4** Modified Halo-tag template synthesis: amplification of Halo-tag sequence using the following primers

F: /5Sp9/A*a*t*tggaagcaaatgacatc

R: /5Sp9/a*t*c*agcttcagttcattctt

/5Sp9/=triethylene glycol; *= phosphorothioate bonds

### **Supplementary Table 3**. Primers used for high throughput sequencing.

| Gene locus | F (5'-3') | R (5'-3') |
|---|---|---|
| *AAVS1-N* | AGACGTGTGCTCTTCCGATCTcgacaaccactacctgagca | CTACACGACGCTCTTCCGATCTccaggatcagtgaaacgcac |
| *AAVS1-Mid* | AGACGTGTGCTCTTCCGATCT gctaccccgaccacatgaa | CTACACGACGCTCTTCCGATCTtccttgaagtcgatgccctt |
| *AAVS-C* | AGACGTGTGCTCTTCCGATCT cttgccagaacctctaaggt | CTACACGACGCTCTTCCGATCTaacttgtggccgtttacgt |

### **Supplementary Table 4**. Primers used for droplet digital PCR (ddPCR).

| Gene locus | 5'-3' |
|---|---|
| PPP1R12C | Wt probe acaatcctaggaagcagggtcagcc |
| AAVS1 | ins probe CACCACCCCGGTGAACAGCTC |
| PPP1R12C-F | ggctgagagctttagaggcg |
| PPP1R12C-R | gacagtcccagttggtaggg |
| AAVS1-F | cttgccagaacctctaaggt |
| AAVS1-R | gctgaacttgtggccgttt |
| 643_HEK3_halo_ddPCR_F | atgcatttgtaggcttgatg |
| 644_HEK3_halo_ddPCR_R | gccaaagacctactactctg |
| 645_HEK3_halo_probe | [6FAM]actgagcacgAattggaagc/3BHQ1/ |

**SUPPLEMENTARY NOTE1:**

We observed that **AAVS1_peg1** tends to generate more indels compared to **AAVS1_peg1B**, likely due to the GA-rich repetitive sequences present in the RTT region of peg1.

**SUPPLEMENTARY NOTE2:**

We designed a pair of pegRNAs targeting the *HEK3* locus to direct 1-kb target deletion and concurrent insertion of two 20-bp flaps homologous to the ends of a linear double-stranded DNA donor carrying a 1.5-kb Halo-tag sequence (**Extended Data Fig. 5B**, left panel). Previous reports show that RNA::triethylene glycol (TEG) modified HDR repair template can improve Cas9-mediated HDR efficiency. Inspired by these findings, we designed a set of primers containing a TEG modification (labeled as 5Sp9) at its 5' end and phosphorothioate bonds at the first three nucleotides. Amplification of the native DNA template using this primer set can incorporate TEG modification and bonds into the 5' ends of the DNA template. Encouragingly, in the presence of the chemically modified template, PE-Cas9 can program 4.71±0.923% desired large deletion and concurrent insertion (**Extended Data Fig. 5B**, right panel). To benchmark it against the existing insertion methods, we include a Cas9-mediated NHEJ-based large insertion approach—HITI—as a reference. PE-Cas9 achieved significantly higher insertion efficiency compared to the Cas9-mediated HITI approach in HEK293T cells.

**SUPPLEMENTARY SEQUENCES**. DsDNA/ssDNA donor sequences and plasmids template sequences

#### 0.8k insertion at *AAVS1* site with AAVS1_peg1B and AAVS1_peg2

FlapA
FlapB
Multi-donor overlap

ATGTCTGGTCAACCACCGCGGTCTCAGTGGTGTACaagcttctgacctcttctcttcctcccacagggcctcgagagatctggcagcgg
agagggcagaggaagtctgctaacatgcggtgacgtcgaggagaatcctggcccagtgagcaagggcgaggagctgttcaccggggtgg
tgcccatcctggtcgagctggacggcgacgtaaacggccacaagttcagcgtgtccggcgagggcgagggcgatgccacctacggcaag
ctgaccctgaagttcatctgcaccaccggcaagctgcccgtgccctggcccaccctcgtgaccaccctgacctacggcgtgcagtgctt
cagccgctaccccgaccacatgaagcagcacgacttcttcaagtccgccatgcccgaaggctacgtccaggagcgcaccatcttcttca
aggacgacggcaactacaagacccgcgccgaggtgaagttcgagggcgacaccctggtgaaccgcatcgagctgaagggcatcgacttc
aaggaggacggcaacatcctggggcacaagctggagtacaactacaacagccacaacgtctatatcatggccgacaagcagaagaacgg
catcaaggtgaacttcaagatccgccacaacatcgaggacggcagcgtgcagctcgccgaccactaccagcagaacacccccatcggcg
acggccccgtgctgctgcccgacaaccactacctgagcacccagtccgccctgagcaaagaccccaacgagaagcgcgatcacatggtc
ctgctggagttcgtgaccgccgccgggatcacTCTCGGCATGGACGAGCTGTACAAGGAATTCTGAG

#### 2.2k insertion at *AAVS1* site with AAVS1_peg1B and AAVS1_peg3

ATGTCTGGTCAACCACCGCGGTCTCAGTGGTGTACaagcttctgacctcttctcttcctcccacagggcctcgagagatctggcagcgg
agagggcagaggaagtctgctaacatgcggtgacgtcgaggagaatcctggcccagtgagcaagggcgaggagctgttcaccggggtgg
tgcccatcctggtcgagctggacggcgacgtaaacggccacaagttcagcgtgtccggcgagggcgagggcgatgccacctacggcaag
ctgaccctgaagttcatctgcaccaccggcaagctgcccgtgccctggcccaccctcgtgaccaccctgacctacggcgtgcagtgctt
cagccgctaccccgaccacatgaagcagcacgacttcttcaagtccgccatgcccgaaggctacgtccaggagcgcaccatcttcttca
aggacgacggcaactacaagacccgcgccgaggtgaagttcgagggcgacaccctggtgaaccgcatcgagctgaagggcatcgacttc
aaggaggacggcaacatcctggggcacaagctggagtacaactacaacagccacaacgtctatatcatggccgacaagcagaagaacgg
catcaaggtgaacttcaagatccgccacaacatcgaggacggcagcgtgcagctcgccgaccactaccagcagaacacccccatcggcg
acggccccgtgctgctgcccgacaaccactacctgagcacccagtccgccctgagcaaagaccccaacgagaagcgcgatcacatggtc
ctgctggagttcgtgaccgccgccgggatcactctcggcatggacgagctgtacaaggaattctgagcccagaacctgagctgcggacc
gactcggtcccactttttcaagttgataacggactagccttattttaacttgctatttctagctctaaaacggatcctctctggctcca
tcggtgtttcgtcctttccacaagatatataaagccaagaaatcgaaatgtatcgtatatgcaaatatgaaggaatcatgggaaatagg
ccctcggtgaagggggcggccgccaccgcggtggagctccagcttttgttccctttagtgagggttaattgcgcgcttggcgtaatcat
ggtcatagctgtttcctgtgtgaaattgttatccgctcacaattccacacaacatacgagccggaagcataaagtgtaaagcctggggt
gcctaatgagtgagctaactcacattaattgcgttgcgctcactgcccgctttccagtcgggaaacctgtcgtgccagctgcattaatg
aatcggccaacgcgcggggagaggcggtttgcgtattgggcgctcttccgcttcctcgctcactgactcgctgcgctcggtcgttcggc
tgcggcgagcggtatcagctcactcaaaggcggtaatacggttatccacagaatcaggggataacgcaggaaagaacatgtgagcaaaa
ggccagcaaaaggccaggaaccgtaaaaaggccgcgttgctggcgtttttccataggctccgcccccctgacgagcatcacaaaaatcg
acgctcaagtcagaggtggcgaaacccgacaggactataaagataccaggcgtttccccctggaagctccctcgtgcgctctcctgttc
cgaccctgccgcttaccggatacctgtccgcctttctcccttcgggaagcgtggcgctttctcatagctcacgctgtaggtatctcagt
tcggtgtaggtcgttcgctccaagctgggctgtgtgcacgaaccccccgttcagcccgaccgctgcgccttatccggtaactatcgtct
tgagtccaacccggtaagacacgacttatcgccactggcagcagccactggtaacaggattagcagagcgaggtatgtaggcggtgcta
cagagttcttgaagtggtggcctaactacggctacactagaagaacagtatttggtatctgcgctctgctgaagccagttaccttcgga
aaaagagttggtagctcttgatccggcaaacaaaccaccgctggtagcggtggtttttttgtttgcaagcagcagattacgcgcagaaa
aaaaggatctcaagaagatcctttgatcttttctacggggtctgacgctcagtggCCGCGGTGGAGCTCCAATTCGCCCTATAGTGAGT
C

#### 4k insertion at *AAVS1* site with AAVS1_peg1B and AAVS1_peg3

ATGTCTGGTCAACCACCGCGGTCTCAGTGGTGTACaagcttctgacctcttctcttcctcccacagggcctcgagagatctggcagcgg
agagggcagaggaagtctgctaacatgcggtgacgtcgaggagaatcctggcccagtgagcaagggcgaggagctgttcaccggggtgg

tgcccatcctggtcgagctggacggcgacgtaaacggccacaagttcagcgtgtccggcgagggcgagggcgatgccacctacggcaag
ctgaccctgaagttcatctgcaccaccggcaagctgcccgtgccctggcccaccctcgtgaccaccctgacctacggcgtgcagtgctt
cagccgctaccccgaccacatgaagcagcacgacttcttcaagtccgccatgcccgaaggctacgtccaggagcgcaccatcttcttca
aggacgacggcaactacaagacccgcgccgaggtgaagttcgagggcgacaccctggtgaaccgcatcgagctgaagggcatcgacttc
aaggaggacggcaacatcctggggcacaagctggagtacaactacaacagccacaacgtctatatcatggccgacaagcagaagaacgg
catcaaggtgaacttcaagatccgccacaacatcgaggacggcagcgtgcagctcgccgaccactaccagcagaacacccccatcggcg
acggccccgtgctgctgcccgacaaccactacctgagcacccagtccgccctgagcaaagaccccaacgagaagcgcgatcacatggtc
ctgctggagttcgtgaccgccgccgggatcactctcggcatggacgagctgtacaaggaattctgagcccagaacctgagctgcggacc
gactcggtcccactttttcaagttgataacggactagccttattttaacttgctatttctagctctaaaacggatcctctctggctcca
tcggtgtttcgtcctttccacaagatatataaagccaagaaatcgaaatgtatcgtatatgcaaatatgaaggaatcatgggaaatagg
ccctcggtgaagggggcggccgccaccgcggtggagctccagcttttgttccctttagtgagggttaattgcgcgcttggcgtaatcat
ggtcatagctgtttcctgtgtgaaattgttatccgctcacaattccacacaacatacgagccggaagcataaagtgtaaagcctggggt
gcctaatgagtgagctaactcacattaattgcgttgcgctcactgcccgctttccagtcgggaaacctgtcgtgccagctgcattaatg
aatcggccaacgcgcggggagaggcggtttgcgtattgggcgctcttccgcttcctcgctcactgactcgctgcgctcggtcgttcggc
tgcggcgagcggtatcagctcactcaaaggcggtaatacggttatccacagaatcaggggataacgcaggaaagaacatgtgagcaaaa
ggccagcaaaaggccaggaaccgtaaaaaggccgcgttgctggcgtttttccataggctccgcccccctgacgagcatcacaaaaatcg
acgctcaagtcagaggtggcgaaacccgacaggactataaagataccaggcgtttccccctggaagctccctcgtgcgctctcctgttc
cgaccctgccgcttaccggatacctgtccgcctttctcccttcgggaagcgtggcgctttctcatagctcacgctgtaggtatctcagt
tcggtgtaggtcgttcgctccaagctgggctgtgtgcacgaaccccccgttcagcccgaccgctgcgccttatccggtaactatcgtct
tgagtccaacccggtaagacacgacttatcgccactggcagcagccactggtaacaggattagcagagcgaggtatgtaggcggtgcta
cagagttcttgaagtggtggcctaactacggctacactagaagaacagtatttggtatctgcgctctgctgaagccagttaccttcgga
aaaagagttggtagctcttgatccggcaaacaaaccaccgctggtagcggtggtttttttgtttgcaagcagcagattacgcgcagaaa
aaaaggatctcaagaagatcctttgatcttttctacggggtctgacgctcagtggaacgaaaactcacgttaagggattttggtcatga
gattatcaaaaaggatcttcacctagatccttttaaattaaaaatgaagttttaaatcaatctaaagtatatatgagtaaacttggtct
gacagttaccaatgcttaatcagtgaggcacctatctcagcgatctgtctatttcgttcatccatagttgcctgactccccgtcgtgta
gataactacgatacgggagggcttaccatctggccccagtgctgcaatgataccgcgagacccacgctcaccggctccagatttatcag
caataaaccagccagccggaagggccgagcgcagaagtggtcctgcaactttatccgcctccatccagtctattaattgttgccgggaa
gctagagtaagtagttcgccagttaatagtttgcgcaacgttgttgccattgctacaggcatcgtggtgtcacgctcgtcgtttggtat
ggcttcattcagctccggttcccaacgatcaaggcgagttacatgatcccccatgttgtgcaaaaaagcggttagctccttcggtcctc
cgatcgttgtcagaagtaagttggccgcagtgttatcactcatggttatggcagcactgcataattctcttactgtcatgccatccgta
agatgcttttctgtgactggtgagtactcaaccaagtcattctgagaatagtgtatgcggcgaccgagttgctcttgcccggcgtcaat
acgggataataccgcgccacatagcagaactttaaaagtgctcatcattggaaaacgttcttcggggcgaaaactctcaaggatcttac
cgctgttgagatccagttcgatgtaacccactcgtgcacccaactgatcttcagcatcttttactttcaccagcgtttctgggtgagca
aaaacaggaaggcaaaatgccgcaaaaaagggaataagggcgacacggaaatgttgaatactcatactcttcctttttcaatattattg
aagcatttatcagggttattgtctcatgagcggatacatatttgaatgtatttagaaaaataaacaaataggggttccgcgcacatttc
cccgaaaagtgccacctaaattgtaagcgttaatattttgttaaaattcgcgttaaatttttgttaaatcagctcattttttaaccaat
aggccgaaatcggcaaaatcccttataaatcaaaagaatagaccgagatagggttgagtgttgttccagtttggaacaagagtccacta
ttaaagaacgtggactccaacgtcaaagggcgaaaaaccgtctatcagggcgatggcccactacgtgaaccatcaccctaatcaagttt
tttggggtcgaggtgccgtaaagcactaaatcggaaccctaaagggagcccccgatttagagcttgacggggaaagccggcgaacgtgg
cgagaaaggaagggaagaaagcgaaaggagcgggcgctagggcgctggcaagtgtagcggtcacgctgcgcgtaaccaccacacccgcc
gcgcttaatgcgccgctacagggcgcgtcccattcgccattcaggctgcgcaactgttgggaagggcgatcggtgcgggcctcttcgct
attacgccagctggcgaaagggggatgtgctgcaaggcgattaagttgggtaacgccagggttttcccagtcacgacgttgtaaaacga
cggccagtgagcgcgcgtaatacgactcactatagggcgaattgggtaccgggccccccctcgaggtcgacggtatcgataagcttgat
gtcgaattcaaaaaaatttctagttggtttaacgcgtaactagatagaaccgcgtcaagagagagagagccccgcggtggagctccaat
tcgccctatagtgagtc

**95 bp ssDNA insertion at *AAVS1* site with AAVS1_peg1B and AAVS1_peg2**

ATGTCTGGTCAACCACCGCGGTCTCAGTGGTGTACcgaTAGGGATAACAGGGTAATgccgtctcggcatggacgagc
tgtacaaggaattctgag

***AAT* 1.9k insertion at *AAVS1* site with AAVS1_peg1 and AAVS1_peg2**

AAGCTTCTGACCTCTTCTCTTCCTCCCACAGGGCCtgtaagtagcctgactagcctagctcgaggttaacgaattccgccccccccc
taacgttactggccgaagccgcttggaataaggccggtgtgcgtttgtctatatgttattttccaccatattgccgtcttttggcaatg
tgagggcccggaaacctggccctgtcttcttgacgagcattcctaggggtctttcccctctcgccaaaggaatgcaaggtctgttgaat
gtcgtgaaggaagcagttcctctggaagcttcttgaagacaaacaacgtctgtagcgaccctttgcaggcagcggaaccccccacctgg
cgacaggtgcctctgcggccaaaagccacgtgtataagatacacctgcaaaggcggcacaaccccagtgccacgttgtgagttggatag
ttgtggaaagagtcaaatggctctcctcaagcgtattcaacaaggggctgaaggatgcccagaaggtaccccattgtatgggatctgat
ctggggcctcggtgcacatgctttacatgtgtttagtcgaggttaaaaaacgtctaggccccccgaaccacggggacgtggttttcctt
tgaaaaacacgatgataatatggccacaaccatgccatcctccgtttcctggggcatcctcctgctggcaggcctgtgctgcctggtcc
ctgtctccctggctgaggatccccagggagatgctgcccagaagacagatacatcccaccatgatcaggatcacccaaccttcaacaag
atcacccccaacctggctgagttcgccttcagcctataccgccagctggcacaccagtccaacagcaccaatatcttcttctccccagt
gagcatcgctacagcctttgcaatgctctccctggggaccaaggctgacactcacgatgaaatcctggagggcctgaatttcaacctca
cggagattccggaggctcagatccatgaaggcttccaggaactcctccgtaccctcaaccagccagacagccagctccagctgaccacc
ggcaatggcctgttcctcagcgagggcctgaagctagtggataagtttttggaggatgttaaaaagttgtaccactcagaagccttcac
tgtcaacttcggggacaccgaagaggccaagaaacagatcaacgattacgtggagaagggtactcaagggaaaattgtggatttggtca
aggagcttgacagagacacagttttgctctggtgaattacatcttctttaaaggcaaatgggagagaccctttgaagtcaaggacacc
gaggaagaggacttccacgtggaccaggtgaccaccgtgaaggtgcctatgatgaagcgtttaggcatgtttaacatccagcactgtaa
gaagctgtccagctgggtgctgctgatgaaatacctgggcaatgccaccgccatcttcttcctgcctgatgaggggaaactacagcacc
tggaaaatgaactcacccacgatatcatcaccaagttcctggaaaatgaagacagaaggtctgccagcttacatttacccaaactgtcc
attactggaacctatgatctgaagagcgtcctgggtcaactgggcatcactaaggtcttcagcaatggggctgacctctccggggtcac
agaggaggcacccctgaagctctccaaggccgtgcataaggctgtgctgaccatcgacgagaaagggactgaagctgctggggccatgt
ttttagaggccatacccatgtctatccccccgaggtcaagttcaacaaaccctttgtcttcttaatgattgaacaaaataccaagtct
cccctcttcatgggaaaagtggtgaatcccacccaaaaagagcagaagctgatcagcgaggaggacctgtagcctgcagggcTCTCGGC
ATGGACGAGCTGTACAAGGAATTCTGAG

**AAVS1-0.8k-two-segments-1**

AAGCTTCTGACCTCTTCTCTTCCTCCCACAGGGCCtcgagagatctggcagcggagagggcagaggaagtctgctaacatgcggtgacg
tcgaggagaatcctggcccagtgagcaagggcgaggagctgttcaccggggtggtgcccatcctggtcgagctggacggcgacgtaaac
ggccacaagttcagcgtgtccggcgagggcgagggcgatgccacctacggcaagctgaccctgaagttcatctgcaccaccggcaagct
gcccgtgccctggcccaccctcgtgaccaccctgacctacggcgtgcagtgcttcagccgctaccccgaccacatgaagcagcacgact
tcttcaagtccgccatgcccgaaggctacgtccaggagcgcaccatCTTCTTCAAGGACGACGGCAACTACAAGAC

**AAVS1-0.8k-two-segments-2**

CTTCTTCAAGGACGACGGCAACTACAAGACccgcgccgaggtgaagttcgagggcgacaccctggtgaaccgcatcgagctgaagggca
tcgacttcaaggaggacggcaacatcctggggcacaagctggagtacaactacaacagccacaacgtctatatcatggccgacaagcag
aagaacggcatcaaggtgaacttcaagatccgccacaacatcgaggacggcagcgtgcagctcgccgaccactaccagcagaacacccc
catcggcgacggccccgtgctgctgcccgacaaccactacctgagcacccagtccgccctgagcaaagaccccaacgagaagcgcgatc
acatggtcctgctggagttcgtgaccgccgccgggatcacTCTCGGCATGGACGAGCTGTACAAGGAATTCTGAG

**AAVS1-AAT-1.9k-Three-segments-1**

AAGCTTCTGACCTCTTCTCTTCCTCCCACAGGGCCtgtaagtagcctgactagcctagctcgaggttaacgaattccgccccccccc
taacgttactggccgaagccgcttggaataaggccggtgtgcgtttgtctatatgttattttccaccatattgccgtcttttggcaatg
tgagggcccggaaacctggccctgtcttcttgacgagcattcctaggggtctttcccctctcgccaaaggaatgcaaggtctgttgaat
gtcgtgaaggaagcagttcctctggaagcttcttgaagacaaacaacgtctgtagcgaccctttgcaggcagcggaaccccccacctgg
cgacaggtgcctctgcggccaaaagccacgtgtataagatacacctgcaaaggcggcacaaccccagtgccacgttgtgagttggatag
ttgtggaaagagtcaaatggctctcctcaagcgtattcaacaaggggctgaaggatgcccagaaggtaccccattgtatgggatctgat
ctggggcctcggtgcacatgctttacatgtgtttagtcgaggttaaaaaacgtctaggccccccgaaccacggggacgtggttttcctt

tgaaaaacacgatgataatatggccacaaccatgccatcctccgtttcctggggcatcctcctgctggcaggcctgtgctgcctggtcc
ctgtCTCCCTGGCTGAGGATCCCCAGGGAGATGC

**AAVS1-AAT-1.9k-Three-segments-2**

CTCCCTGGCTGAGGATCCCCAGGGAGATGCtgcccagaagacagatacatcccaccatgatcaggatcacccaaccttcaacaagatca
cccccaacctggctgagttcgccttcagcctataccgccagctggcacaccagtccaacagcaccaatatcttcttctccccagtgagc
atcgctacagcctttgcaatgctctccctggggaccaaggctgacactcacgatgaaatcctggagggcctgaatttcaacctcacgga
gattccggaggctcagatccatgaaggcttccaggaactcctccgtaccctcaaccagccagacagccagctccagctgaccaccggca
atggcctgttcctcagcgagggcctgaagctagtggataagtttttggaggatgttaaaaagttgtaccactcagaagccttcactgtc
aacttcggggacaccgaagaggccaagaaacagatcaacgattacgtggagaagggtactcaagggaaaattgtggatttggtcaagga
gcttgacagagacacagtttttgctctggtgaattacatcttctttaaaggcaaatgggagagaccctttgaagtcaaggacaccgagg
aagaggacttccacgtggaccaggtgaccaccgtgaaggtgcctatgatgaagcgtttaggcatgtttaaCATCCAGCACTGTAAGAAG
CTGTCCAGCTG

**AAVS1-AAT-1.9k-Three-segments-3**

CATCCAGCACTGTAAGAAGCTGTCCAGCTGggtgctgctgatgaaatacctgggcaatgccaccgccatcttcttcctgcctgatgaggggaaactac
agcacctggaaaatgaactcacccacgatatcatcaccaagttcctggaaaatgaagacagaaggtctgccagcttacatttacccaaactgtccatt
actggaacctatgatctgaagagcgtcctgggtcaactgggcatcactaaggtcttcagcaatggggctgacctctccggggtcacagaggaggcacc
cctgaagctctccaaggccgtgcataaggctgtgctgaccatcgacgagaaagggactgaagctgctggggccatgtttttagaggccatacccatgt
ctatcccccccgaggtcaagttcaacaaaccctttgtcttcttaatgattgaacaaaataccaagtctcccctcttcatgggaaaagtggtgaatccc
acccaaaaagagcagaagctgatcagcgaggaggacctgtagcctgcagggcTCTCGGCATGGACGAGCTGTACAAGGAATTCTGAG

***TRAC*-0.8k**

ATGTCTGGTCAACCACCGCGGTCTCAGTGGTGTACaagcttctgacctcttctcttcctcccacagATggcctcgagagatctggcagc
ggagagggcagaggaagtctgctaacatgcggtgacgtcgaggagaatcctggcccagtgagcaagggcgaggagctgttcaccggggt
ggtgcccatcctggtcgagctggacggcgacgtaaacggccacaagttcagcgtgtccggcgagggcgagggcgatgccacctacggca
agctgaccctgaagttcatctgcaccaccggcaagctgcccgtgccctggcccaccctcgtgaccaccctgacctacggcgtgcagtgc
ttcagccgctaccccgaccacatgaagcagcacgacttcttcaagtccgccatgcccgaaggctacgtccaggagcgcaccatcttctt
caaggacgacggcaactacaagacccgcgccgaggtgaagttcgagggcgacaccctggtgaaccgcatcgagctgaagggcatcgact
tcaaggaggacggcaacatcctggggcacaagctggagtacaactacaacagccacaacgtctatatcatggccgacaagcagaagaac
ggcatcaaggtgaacttcaagatccgccacaacatcgaggacggcagcgtgcagctcgccgaccactaccagcagaacacccccatcgg
cgacggccccgtgctgctgcccgacaaccactacctgagcacccagtccgccctgagcaaagaccccaacgagaagcgcgatcacatgg
tcctgctggagttcgtgaccgccgccgggatcacTCTCGGCATGGACGAGCTGTACAAGGAATTCTGAG

***TRAC*-2.4k**

ATGTCTGGTCAACCACCGCGGTCTCAGTGGTGTACgactacaaggatcatgacggagactataaggatcacgatattgattacaaagat
gacgacgacaaagacatccagatgacacagactacatcctccctgtctgcctctctgggagacagagtcaccatcagttgcagggcaag
tcaggacatctctaagtatttgaattggtatcagcagaaaccagatggaactgttaaactcctgatctaccatacatcaagattacact
caggagtcccatcaaggttcagtggcagtgggtctggaacagattattctctcaccattagcaacctggagcaagaagatattgccact
tacttttgccaacagggtaatacgcttccgtacacgttcggaggggggactaagttggaaataacaggctccacctctggatccggcaa
gcccggatctggcgagggatccaccaagggcgaggtgaaactgcaggagtcaggacctggcctggtggcgccctcacagagcctgtccg
tcacatgcactgtctcaggggtctcattacccgactatggtgtaagctggattcgccagcctccacgaaagggtctggagtggctggga
gtaatatggggtagtgaaaccacatactataattcagctctcaaatccagactgaccatcatcaaggacaactccaagagccaagtttt
cttaaaaatgaacagtctgcaaactgatgacacagccatttactactgtgccaaacattattactacggtggtagctatgctatggact
actggggtcaaggaacctcagtcaccgtctcctcagcggccgcaggtaccaccacaacgcccgctcctcggccaccgacgccagcgcca
actattgcgagtcagcctctcagtctgcgacctgaggcttgtcgaccagcagccggaggcgcagtgcacacgagggggctggacttcgc
ctgtgatagaagacctccttctaagcccttttgggtgctggtggtggttggtggagtcctggcttgctatagcttgctagtaacagtgg
cctttattattttctgggtgaggagtaagaggagcaggctcctgcacagtgactacatgaacatgactcccaggcggcccggacccacc
cgcaagcattaccagccctatgccccaccacgcgacttcgcagcctatcgctccgctagcctgagagtgaagttcagcaggagcgcaga
cgcccccgcgtaccagcagggccagaaccagctctataacgagctcaatctaggacgaagagaggagtacgatgttttggacaagagac
gtggccgggaccctgagatgggggaaagccgcagagaaggaagaaccctcaggaaggcctgtacaatgaactgcagaaagataagatg

gcggaggcctacagtgagattgggatgaaaggcgagcgccggaggggcaaggggcacgatggcctttaccagggtctcagtacagccac
caaggacacctacgacgcccttcacatgcaggccctgcccctcgcgctagcgccacgaacttctctctgttaaagcaagcaggcgacg
tggaagaaaaccccggtcccgtgagcaagggcgaggagctgttcaccggggtggtgcccatcctggtcgagctggacggcgacgtaaac
ggccacaagttcagcgtgtccggcgagggcgagggcgatgccacctacggcaagctgaccctgaagttcatctgcaccaccggcaagct
gcccgtgccctggcccaccctcgtgaccaccctgacctacggcgtgcagtgcttcagccgctaccccgaccacatgaagcagcacgact
tcttcaagtccgccatgcccgaaggctacgtccaggagcgcaccatcttcttcaaggacgacggcaactacaagacccgcgccgaggtg
aagttcgagggcgacaccctggtgaaccgcatcgagctgaagggcatcgacttcaaggaggacggcaacatcctggggcacaagctgga
gtacaactacaacagccacaacgtctatatcatggccgacaagcagaagaacggcatcaaggtgaacttcaagatccgccacaacatcg
aggacggcagcgtgcagctcgccgaccactaccagcagaacacccccatcggcgacggccccgtgctgctgcccgacaaccactacctg
agcacccagtccgccctgagcaaagaccccaacgagaagcgcgatcacatggtcctgctggagttcgtgaccgccgccgggatcactct
cggcatggacgagctgtacaagtaacgcgCCGCGGTGGAGCTCCAATTCGCCCTATAGTGAGTC

***DMD*-11.6k**

ATGTCTGGTCAACCACCGCGGTCTCAGTGGTGTACagaattccgccaccatgctttggtgggaagaagtagaggactgttatgaaagag
aagatgttcaaaagaaaacattcacaaaatgggtaaatgcacaattttctaagtttgggaagcagcatattgagaacctcttcagtgac
ctacaggatgggaggcgcctcctagacctcctcgaaggcctgacagggcaaaaactggtatgtatcaagcttacaagacagctttaagg
agaccaatagaaactggggcatgtggagacagagaagactcttgggcctcgaggcttcatttttctttttctttttcagccaaaagaa
aaaggatccacaagagttcatgccctgaacaatgtcaacaaggcactgcgggttttgcagaacaataatgttgatttagtgaatattgg
aagtactgacatcgtagatggaaatcataaactgactcttggtttgatttggaatataatcctccactggcaggtcaaaaatgtaatga
aaaatatcatggctggattgcaacaaaccaacagtgaaaagattctcctgagctgggtccgacaatcaactcgtaattatccacaggtt
aatgtaatcaacttcaccaccagctggtctgatggcctggctttgaatgctctcatccatagtcataggccagacctatttgactggaa
tagtgtggtttgccagcagtcagccacacaacgactggaacatgcattcaacatcgccagatatcaattaggcatagagaaactactcg
atcctgaagatgttgataccacctatccagataagaagtccatcttaatgtacatcacatcactcttccaagttttgcctcaacaagtg
agcatcgaagccatccaggaagtggaaatgttgccaaggccacctaaagtgactaaagaagaacattttcagttacatcatcaaatgca
ctattctcaacagatcacggtcagtctagcacagggatatgagagaacttcttcccctaagcctcgattcaagagctatgcctacacac
aggctgcttatgtcaccacctctgaccctacacggagcccatttccttcacagcatttggaagctcctgaagacaagtcatttggcagt
tcattgatggagagtgaagtaaacctggaccgttatcaaacagctttagaagaagtattatcgtggcttctttctgctgaggacacatt
gcaagcacaaggagagatttctaatgatgtggaagtggtgaaagaccagtttcatactcatgaggggtacatgatggatttgacagccc
atcagggccgggttggtaatattctacaattgggaagtaagctgattggaacaggaaaattatcagaagatgaagaaactgaagtacaa
gagcagatgaatctcctaaattcaagatgggaatgcctcagggtagctagcatggaaaaacaaagcaatttacatagagttttaatgga
tctccagaatcagaaactgaaagagttgaatgactggctaacaaaaacagaagaaagaacaaggaaaatggaggaagagcctcttggac
ctgatcttgaagacctaaaacgccaagtacaacaacataaggtgcttcaagaagatctagaacaagaacaagtcagggtcaattctctc
actcacatggtggtggtagttgatgaatctagtggagatcacgcaactgctgctttggaagaacaacttaaggtattgggagatcgatg
ggcaaacatctgtagatggacagaagaccgctgggttcttttacaagacatccttctcaaatggcaacgtcttactgaagaacagtgcc
tttttagtgcatggctttcagaaaaagaagatgcagtgaacaagattcacacaactggctttaaagatcaaaatgaaatgttatcaagt
cttcaaaaactggccgttttaaaagcggatctagaaaagaaaaagcaatccatgggcaaactgtattcactcaaacaagatcttctttc
aacactgaagaataagtcagtgacccagaagacggaagcatggctggataactttgcccggtgttgggataatttagtccaaaaacttg
aaaagagtacagcacagatttcacaggctgtcaccaccactcagccatcactaacacagacaactgtaatggaaacagtaactacggtg
accacaagggaacagatcctggtaaagcatgctcaagaggaacttccaccaccacctccccaaaagaagaggcagattactgtggattc
tgaaattaggaaaaggttggatgttgatataactgaacttcacagctggattactcgctcagaagctgtgttgcagagtcctgaatttg
caatctttcggaaggaaggcaacttctcagacttaaaagaaaaagtcaatgccatagagcgagaaaaagctgagaagttcagaaaactg
caagatgccagcagatcagctcaggccctggtggaacagatggtgaatgagggtgttaatgcagatagcatcaaacaagcctcagaaca
actgaacagccggtggatcgaattctgccagttgctaagtgagagacttaactggctggagtatcagaacaacatcatcgctttctata
atcagctacaacaattggagcagatgacaactactgctgaaaactggttgaaaatccaacccaccaccccatcagagccaacagcaatt
aaaagtcagttaaaaatttgtaaggatgaagtcaaccggctatcaggtcttcaacctcaaattgaacgattaaaaattcaaagcatagc
cctgaaagagaaaggacaaggacccatgttcctggatgcagactttgtggcctttacaaatcattttaagcaagtcttttctgatgtgc
aggccagagagaaagagctacagacaatttttgacactttgccaccaatgcgctatcaggagaccatgagtgccatcaggacatgggtc
cagcagtcagaaaccaaactctccatacctcaacttagtgtcaccgactatgaaatcatggagcagagactcggggaattgcaggcttt
acaaagttctctgcaagagcaacaaagtggcctatactatctcagcaccactgtgaaagagatgtcgaagaaagcgccctctgaaatta
gccggaaatatcaatcagaatttgaagaaattgagggacgctggaagaagctctcctcccagctggttgagcattgtcaaaagctagag
gagcaaatgaataaactccgaaaaattcagaatcacatacaaaccctgaagaaatggatggctgaagttgatgtttttctgaaggagga
atggcctgcccttggggattcagaaattctaaaaaagcagctgaaacagtgcagacttttagtcagtgatattcagacaattcagccca
gtctaaacagtgtcaatgaaggtgggcagaagataaagaatgaagcagagccagagtttgcttcgagacttgagacagaactcaaagaa
cttaacactcagtgggatcacatgtgccaacaggtctatgccagaaaggaggccttgaagggaggtttggagaaaactgtaagcctcca

gaaagatctatcagagatgcacgaatggatgacacaagctgaagaagagtatcttgagagagattttgaatataaaactccagatgaat
tacagaaagcagttgaagagatgaagagagctaaagaagaggcccaacaaaaagaagcgaaagtgaaactccttactgagtctgtaaat
agtgtcatagctcaagctccacctgtagcacaagaggccttaaaaaaggaacttgaaactctaaccaccaactaccagtggctctgcac
taggctgaatgggaaatgcaagactttggaagaagtttgggcatgttggcatgagttattgtcatacttggagaaagcaaacaagtggc
taaatgaagtagaatttaaacttaaaaccactgaaaacattcctggcggagctgaggaaatctctgaggtgctagattcacttgaaaat
ttgatgcgacattcagaggataacccaaatcagattcgcatattggcacagaccctaacagatggcggagtcatggatgagctaatcaa
tgaggaacttgagacatttaattctcgttggagggaactacatgaagaggctgtaaggaggcaaaagttgcttgaacagagcatccagt
ctgcccaggagactgaaaaatccttacacttaatccaggagtccctcacattcattgacaagcagttggcagcttatattgcagacaag
gtggacgcagctcaaatgcctcaggaagcccagaaaatccaatctgatttgacaagtcatgagatcagtttagaagaaatgaagaaaca
taatcaggggaaggaggctgcccaaagagtcctgtctcagattgatgttgcacagaaaaaattacaagatgtctccatgaagtttcgat
tattccagaaaccagccaattttgagcagcgtctacaagaaagtaagatgattttagatgaagtgaagatgcacttgcctgcattggaa
acaaagagtgtggaacaggaagtagtacagtcacagctaaatcattgtgtgaacttgtataaaagtctgagtgaagtgaagtctgaagt
ggaaatggtgataaagactggacgtcagattgtacagaaaaagcagacggaaaatcccaaagaacttgatgaaagagtaacagctttga
aattgcattataatgagctgggagcaaaggtaacagaaagaaagcaacagttggagaaatgcttgaaattgtcccgtaagatgcgaaag
gaaatgaatgtcttgacagaatggctggcagctacagatatggaattgacaaagagatcagcagttgaaggaatgcctagtaatttgga
ttctgaagttgcctggggaaaggctactcaaaaagagattgagaaacagaaggtgcacctgaagagtatcacagaggtaggagaggcct
tgaaaacagttttgggcaagaaggagacgttggtggaagataaactcagtcttctgaatagtaattggatagctgtcacctcccgagca
gaagagtggttaaatcttttgttggaataccagaaacacatggaaacttttgaccagaatgtggaccacatcacaaagtggatcattca
ggctgacacacttttggatgaatcagagaaaaagaaaccccagcaaaaagaagacgtgcttaagcgtttaaaggcagaactgaatgaca
tacgcccaaaggtggactctacacgtgaccaagcagcaaacttgatggcaaaccacggtgaccactgcaggaaattagtagagccccaa
atctcagagctcaaccatcgatttgcagccatttcacacagaattaagactggaaaggcctccattcctttgaaggaattggagcagtt
taactcagatatacaaaaattgcttgaaccactggaggctgaaattcagcaggggggtgaatctgaaagaggaagacttcaataaagata
tgaatgaagacaatgagggtactgtaaaagaattgttgcaaagaggagacaacttacaacaaagaatcacagatgagagaaagcgagag
gaaataaagataaaacagcagctgttacagacaaaacataatgctctcaaggatttgaggtctcaaagaagaaaaaaggctctagaaat
ttctcatcagtggtatcagtacaagaggcaggctgatgatctcctgaaatgcttggatgacattgaaaaaaaattagccagcctacctg
agcccagagatgaaaggaaaataaaggaaattgatcgggaattgcagaagaagaaagaggagctgaatgcagtgcgtaggcaagctgag
ggcttgtctgaggatggggccgcaatggcagtggagccaactcagatccagctcagcaagcgctggcgggaaattgagagcaaatttgc
tcagtttcgaagactcaactttgcacaaattcacactgtccgtgaagaaacgatgatggtgatgactgaagacatgcctttggaaattt
cttatgtgccttctacttatttgactgaaatcactcatgtctcacaagccctattagaagtggaacaacttctcaatgctcctgacctc
tgtgctaaggactttgaagatctctttaagcaagaggagtctctgaagaatataaaagatagtctacaacaaagctcaggtcggattga
cattattcatagcaagaagacagcagcattgcaaagtgcaacgcctgtggaaagggtgaagctacaggaagctctctcccagcttgatt
tccaatgggaaaaagttaacaaaatgtacaaggaccgacaagggcgatttgacagatctgttgagaaatggcggcgttttcattatgat
ataaagatatttaatcagtggctaacagaagctgaacagtttctcagaaagacacaaattcctgagaattgggaacatgctaaatacaa
atggtatcttaaggaactccaggatggcattgggcagcggcaaactgttgtcagaacattgaatgcaactggggaagaaataattcagc
aatcctcaaaaacagatgccagtattctacaggaaaaattgggaagcctgaatctgcggtggcaggaggtctgcaaacagctgtcagac
agaaaaaagaggctagaagaacaaaagaatatcttgtcagaatttcaaagagatttaaatgaatttgttttatggttggaggaagcaga
taacattgctagtatcccacttgaacctggaaaagagcagcaactaaaagaaaagcttgagcaagtcaagttactggtggaagagttgc
cccctgcgccagggccgaattctcaaacaattaaatgaaactggaggacccgtgcttgtaagtgctcccataagcccagaagagcaagat
aaacttgaaaataagctcaagcagacaaatctccagtggataaaggtttccagagctttacctgagaaacaaggagaaattgaagctca
aataaaagaccttgggcagcttgaaaaaaagcttgaagaccttgaagagcagttaaatcatctgctgctgtggttatctcctattagga
atcagttggaaatttataaccaaccaaaccaagaaggaccatttgacgttaaggaaactgaaatagcagttcaagctaaacaaccggat
gtggaagagattttgtctaaagggcagcatttgtacaaggaaaaaccagccactcagccagtgaagaggaagttagaagatctgagctc
tgagtggaaggcggtaaaccgtttacttcaagagctgagggcaaagcagcctgacctagctcctggactgaccactattggagcctctc
ctactcagactgttactctggtgacacaacctgtggttactaaggaaactgccatctccaaactagaaatgccatcttccttgatgttg
gaggtacctgctctggcagatttcaaccgggcttggacagaacttaccgactggctttctctgcttgatcaagttataaaatcacagag
ggtgatggtgggtgaccttgaggatatcaacgagatgatcatcaagcagaaggcgacaatgcaggatttggaacagaggcgtccccagt
tggaagaactcattaccgctgcccaaaatttgaaaaacaagaccagcaatcaagaggctagaacaatcattacggatcgaattgaaaga
attcagaatcagtgggatgaagtacaagaacaccttcagaaccggaggcaacagttgaatgaaatgttaaaggattcaacacaatggct
ggaagctaaggaagaagctgagcaggtcttaggacaggccagagccaagcttgagtcatggaaggagggtccctatacagtagatgcaa
tccaaaagaaaatcacagaaaccaagcagttggccaaagacctccgccagtggcagacaaatgtagatgtggcaaatgacttggccctg
aaacttctccgggattattctgcagatgataccagaaaagtccacatgataacagagaatatcaatgcctcttggagaagcattcataa
aagggtgagtgagcgagaggctgctttggaagaaactcatagattactgcaacagttccccctggacctggaaaagtttcttgcctggc
ttacagaagctgaaacaactgccaatgtcctacaggatgctacccgtaaggaaaggctcctagaagactccaagggagtaaaagagctg
atgaaacaatggcaagacctccaaggtgaaattgaagctcacacagatgtttatcacaacctgggtgaaaacagccaaaaaatcctgag
atccctggaaggttccgatgatgcagtcctgttacaaagacgtttggataacatgaacttcaagtggagtgaacttcggaaaaagtctc

tcaacattaggtcccatttggaagccagttctgaccagtggaagcgtctgcacctttctctgcaggaacttctggtgtggctacagctg
aaagatgatgaattaagccggcaggcacctattggaggcgactttccagcagttcagaagcagaacgatgtacatagggccttcaagag
ggaattgaaaactaaagaacctgtaatcatgagtattcttgagactgtacgaatatttctgacagagcagcctttggaaggactagaga
aaccctaccaggagcccagagagctgcctcctgaggagagagcccagaatgtcactcggcttctacgaaagcaggctgaggaggtcaat
actgagtgggaaaaattgaacctgcactccgctgactggcagagaaaaatagatgagacccttgaaagactccaggaacttcaagaggc
cacggatgagctggacctcaagctgcgccaagctgaggtgatcaagggatcctggcagcccgtgggcgatctcctcattgactctctcc
aagatcacctcgagaaagtcaaggcacttcgaggagaaattgcgcctctgaaagagaacgtgagccacgtcaatgaccttgctcgccag
cttaccactttgggcattcagctctcaccgtataacctcagcactctggaagacctgaacaccagatggaagcttctgcaggtggccgt
cgaggaccgagtcaggcagctgcatgaagcccacagggactttggtccagcatctcagcactttctttccacgtctgtccagggtccct
gggagagagccatctcgccaaacaaagtgccctactatatcaaccacgagactcaaacaacttgctgggaccatcccaaaatgacagag
ctctaccagtctttagctgacctgaataatgtcagattctcagcttataggactgccatgaaactccgaagactgcagaaggccctttg
cttggatctcttgagcctgtcagctgcatgtgatgccttggaccagcacaacctcaagcaaaatgaccagcccatggatatcctgcaga
ttattaattgtttgaccactatttatgaccgcctggagcaagagcacaacaatttggtcaacgtccctctctgcgtggatatgtgtctg
aactggctgctgaatgtttatgatacgggacgaacagggaggatccgtgtcctgtcttttaaaactggcatcatttccctgtgtaaagc
acatttggaagacaagtacagataccttttcaagcaagtggcaagttcaacaggattttgtgaccagcgcaggctgggcctccttctgc
atgattctatccaaattccaagacagttgggtgaagttgcatcctttgggggcagtaacattgagccaagtgtccggagctgcttccaa
tttgctaataataagccagagatcgaagcggccctcttcctagactggatgagactggaaccccagtccatggtgtggctgcccgtcct
gcacagagtggctgctgcagaaactgccaagcatcaggccaaatgtaacatctgcaaagagtgtccaatcattggattcaggtacagga
gtctaaagcactttaattatgacatctgccaaagctgctttttttctggtcgagttgcaaaaggccataaaatgcactatcccatggtg
gaatattgcactccgactacatcaggagaagatgttcgagactttgccaaggtactaaaaaacaaatttcgaaccaaaaggtattttgc
gaagcatccccgaatgggctacctgccagtgcagactgtcttagagggggacaacatggaaactcccgttactctgatcaacttctggc
cagtagattctgcgcctgcctcgtcccctcagctttcacacgatgatactcattcacgcattgaacattatgctagcaggctagcagaa
atggaaaacagcaatggatcttatctaaatgatagcatctctcctaatgagagcatagatgatgaacatttgttaatccagcattactg
ccaaagtttgaaccaggactccccccctgagccagcctcgtagtcctgcccagatcttgatttccttagagagtgaggaaagaggggagc
tagagagaatcctagcagatcttgaggaagaaaacaggaatctgcaagcagaatatgaccgtctaaagcagcagcacgaacataaaggc
ctgtccccactgccgtcccctcctgaaatgatgcccacctctccccagagtccccgggatgctgagctcattgctgaggccaagctact
gcgtcaacacaaaggccgcctggaagccaggatgcaaatcctggaagaccacaataaacagctggagtcacagttacacaggctaaggc
agctgctggagcaaccccaggcagaggccaaagtgaatggcacaacggtgtcctctccttctacctctctacagaggtccgacagcagt
cagcctatgctgctccgagtggttggcagtcaaacttcggactccatgggtaagtagagggctgggctgggctgtggggggtgtggggt
gcgggactgggcagtctgggagtccctctcaccacttttcttacctttctaggtgaggaagatcttctcagtcctccccaggacacaag
cacagggttagaggaggtgatggagcaactcaacaactccttccctagttcaagaggaagaaataccccctggaaagccaatgagagagg
acacaatgggatccggtgccTCTCGGCATGGACGAGCTGTACAAGGAATTCTGAG

#### *ACTB* GFP insertion with ACTB_peg1 and ACTB_peg2

GCCACCATGGTGAGCAAGGGCGAGGAGCTGTTCACcggggtggtgcccatcctggtcgagctggacggcgacgtaaacggccacaagtt
cagcgtgtccggcgagggcgagggcgatgccacctacggcaagctgaccctgaagttcatctgcaccaccggcaagctgcccgtgccct
ggcccaccctcgtgaccaccctgacctacggcgtgcagtgcttcagccgctaccccgaccacatgaagcagcacgacttcttcaagtcc
gccatgcccgaaggctacgtccaggagcgcaccatcttcttcaaggacgacggcaactacaagacccgcgccgaggtgaagttcgaggg
cgacaccctggtgaaccgcatcgagctgaagggcatcgacttcaaggaggacggcaacatcctggggcacaagctggagtacaactaca
acagccacaacgtctatatcatggccgacaagcagaagaacggcatcaaggtgaacttcaagatccgccacaacatcgaggacggcagc
gtgcagctcgccgaccactaccagcagaacacccccatcggcgacggccccgtgctgctgcccgacaaccactacctgagcacccagtc
cgccctgagcaaagaccccaacgagaagcgcgatcacatggtcctgctggagttcgtgaccgccgccgggatcactctcggcatggacg
agctgtacaagggatctggtgccacgaatttctcgctactcaagcaggccggtgatgtcgaggaaaaccctggtcctatgTGCAAGGCC
GGATTTGCAGGCGATGATGCTCCACG

#### *ACTB* GFP insertion with ACTB_peg1 and ACTB_peg3

GCCACCATGGTGAGCAAGGGCGAGGAGCTGTTCACcggggtggtgcccatcctggtcgagctggacggcgacgtaaacggccacaagtt
cagcgtgtccggcgagggcgagggcgatgccacctacggcaagctgaccctgaagttcatctgcaccaccggcaagctgcccgtgccct
ggcccaccctcgtgaccaccctgacctacggcgtgcagtgcttcagccgctaccccgaccacatgaagcagcacgacttcttcaagtcc
gccatgcccgaaggctacgtccaggagcgcaccatcttcttcaaggacgacggcaactacaagacccgcgccgaggtgaagttcgaggg
cgacaccctggtgaaccgcatcgagctgaagggcatcgacttcaaggaggacggcaacatcctggggcacaagctggagtacaactaca
acagccacaacgtctatatcatggccgacaagcagaagaacggcatcaaggtgaacttcaagatccgccacaacatcgaggacggcagc
gtgcagctcgccgaccactaccagcagaacacccccatcggcgacggccccgtgctgctgcccgacaaccactacctgagcacccagtc

cgccctgagcaaagaccccaacgagaagcgcgatcacatggtcctgctggagttcgtgaccgccgccgggatcactctcggcatggacg
agctgtacaagggatctggtgccacgaatttctcgctactcaagcaggccggtgatgtcgaggaaaaccctggtcctatggacgacgac
atagctgcattagtagtggataatggttctggaatgTGTAAAGCTGGTTTTGCAGGTGATGACGCTCCTCG

#### Plasmid template sequences for 0.8/2.2/4k insertions at *AAVS1*

AAAAAAACCACCGCTACCAGCGGTGGTTTGTTTGCCGGATCAAGAGCTACCAACTCTTTTTCCGAAGGTAACTGGCTTCAGCAGAGCGC
AGATACCAAATACTGTTCTTCTAGTGTAGCCGTAGTTAGGCCACCACTTCAAGAACTCTGTAGCACCGCCTACATACCTCGCTCTGCTA
ATCCTGTTACCAGTGGCTGCTGCCAGTGGCGATAAGTCGTGTCTTACCGGGTTGGACTCAAGACGATAGTTACCGGATAAGGCGCAGCG
GTCGGGCTGAACGGGGGGTTCGTGCACACAGCCCAGCTTGGAGCGAACGACCTACACCGAACTGAGATACCTACAGCGTGAGCTATGAG
AAAGCGCCACGCTTCCCGAAGGGAGAAAGGCGGACAGGTATCCGGTAAGCGGCAGGGTCGGAACAGGAGAGCGCACGAGGGAGCTTCCA
GGGGGAAACGCCTGGTATCTTTATAGTCCTGTCGGGTTTCGCCACCTCTGACTTGAGCGTCGATTTTTGTGATGCTCGTCAGGGGGGCG
GAGCCTATGGAAAAACGCCAGCAACGCGGCCTTTTTACGGTTCCTGGCCTTTTGCTGGCCTTTTGCTCACATGTTCTTTCCTGCGTTAT
CCCCTGATTCTGTGGATAACCGTATTACCGCCTTTGAGTGAGCTGATACCGCTCGCCGCAGCCGAACGACCGAGCGCAGCGAGTCAGTG
AGCGAGGAAGCGGAAGAGCGCCCAATACGCAAACCGCCTCTCCCCGCGCGTTGGCCGATTCATTAATGCAGCTGGCACGACAGGTTTCC
CGACTGGAAAGCGGGCAGTGAGCGCAACGCAATTAATGTGAGTTAGCTCACTCATTAGGCACCCCAGGCTTTACACTTTATGCTTCCGG
CTCGTATGTTGTGTGGAATTGTGAGCGGATAACAATTTCACACAGGAAACAGCTATGACCATGATTACGCCAAGCGCGCAATTAACCCT
CACTAAAGGGAACAAAAGCTGGAGCTCCACCGCGGTGGCGGCCGCCCCCTTCACCGAGGGCCTATTTCCGAATTAATTTGACTGTAAAC
ACAAAGATATGGAAAGGACGAAACACCGATGGAGCCAGAGAGGATCCGTTTTAGAGCTAGAAATAGCAAGTTAAAATAAGGCTAGTCCG
TTATCAACTTGAAAAAGTGGGACCGAGTCGGTCCGCAGCTCAGGTTCTGGGCTCAGAATTCCTTGTACAGCTCGTCCATGCCGAGAGTG
ATCCCGGCGGCGGTCACGAACTCCAGCAGGACCATGTGATCGCGCTTCTCGTTGGGGTCTTTGCTCAGGGCGGACTGGGTGCTCAGGTA
GTGGTTGTCGGGCAGCAGCACGGGGCCGTCGCCGATGGGGGTGTTCTGCTGGTAGTGGTCGGCGAGCTGCACGCTGCCGTCCTCGATGT
TGTGGCGGATCTTGAAGTTCACCTTGATGCCGTTCTTCTGCTTGTCGGCCATGATATAGACGTTGTGGCTGTTGTAGTTGTACTCCAGC
TTGTGCCCCAGGATGTTGCCGTCCTCCTTGAAGTCGATGCCCTTCAGCTCGATGCGGTTCACCAGGGTGTCGCCCTCGAACTTCACCTC
GGCGCGGGTCTTGTAGTTGCCGTCGTCCTTGAAGAAGATGGTGCGCTCCTGGACGTAGCCTTCGGGCATGGCGGACTTGAAGAAGTCGT
GCTGCTTCATGTGGTCGGGGTAGCGGCTGAAGCACTGCACGCCGTAGGTCAGGGTGGTCACGAGGGTGGGCCAGGGCACGGGCAGCTTG
CCGGTGGTGCAGATGAACTTCAGGGTCAGCTTGCCGTAGGTGGCATCGCCCTCGCCCTCGCCGGACACGCTGAACTTGTGGCCGTTTAC
GTCGCCGTCCAGCTCGACCAGGATGGGCACCACCCCGGTGAACAGCTCCTCGCCCTTGCTCACTGGGCCAGGATTCTCCTCGACGTCAC
CGCATGTTAGCAGACTTCCTCTGCCCTCTCCGCTGCCAGATCTCTCGAGGCCCTGTGGGAGGAAGAGAAGAGGTCAGAAGCTTTCCTCT
CTGGCTCTCTCTCTCTTGACGCGGTTCTATCTAGTTACGCGTTAAACCAACTAGAAATTTTTTTGAATTCGACATCAAGCTTATCGATA
CCGTCGACCTCGAGGGGGGGCCCGGTACCCAATTCGCCCTATAGTGAGTCGTATTACGCGCGCTCACTGGCCGTCGTTTTACAACGTCG
TGACTGGGAAAACCCTGGCGTTACCCAACTTAATCGCCTTGCAGCACATCCCCCTTTCGCCAGCTGGCGTAATAGCGAAGAGGCCCGCA
CCGATCGCCCTTCCCAACAGTTGCGCAGCCTGAATGGCGAATGGACGCGCCCTGTAGCGGCGCATTAAGCGCGGCGGGTGTGGTGGTT
ACGCGCAGCGTGACCGCTACACTTGCCAGCGCCCTAGCGCCCGCTCCTTTCGCTTTCTTCCCTTCCTTTCTCGCCACGTTCGCCGGCTT
TCCCCGTCAAGCTCTAAATCGGGGGCTCCCTTTAGGGTTCCGATTTAGTGCTTTACGGCACCTCGACCCCAAAAAACTTGATTAGGGTG
ATGGTTCACGTAGTGGGCCATCGCCCTGATAGACGGTTTTTCGCCCTTTGACGTTGGAGTCCACGTTCTTTAATAGTGGACTCTTGTTC
CAAACTGGAACAACACTCAACCCTATCTCGGTCTATTCTTTTGATTTATAAGGGATTTTGCCGATTTCGGCCTATTGGTTAAAAAATGA
GCTGATTTAACAAAAATTTAACGCGAATTTTAACAAAATATTAACGCTTACAATTTAGGTGGCACTTTTCGGGGAAATGTGCGCGGAAC
CCCTATTTGTTTATTTTTCTAAATACATTCAAATATGTATCCGCTCATGAGACAATAACCCTGATAAATGCTTCAATAATATTGAAAAA
GGAAGAGTATGAGTATTCAACATTTCCGTGTCGCCCTTATTCCCTTTTTTGCGGCATTTTGCCTTCCTGTTTTTGCTCACCCAGAAACG
CTGGTGAAAGTAAAAGATGCTGAAGATCAGTTGGGTGCACGAGTGGGTTACATCGAACTGGATCTCAACAGCGGTAAGATCCTTGAGAG
TTTTCGCCCCGAAGAACGTTTTCCAATGATGAGCACTTTTAAAGTTCTGCTATGTGGCGCGGTATTATCCCGTATTGACGCCGGGCAAG
AGCAACTCGGTCGCCGCATACACTATTCTCAGAATGACTTGGTTGAGTACTCACCAGTCACAGAAAAGCATCTTACGGATGGCATGACA
GTAAGAGAATTATGCAGTGCTGCCATAACCATGAGTGATAACACTGCGGCCAACTTACTTCTGACAACGATCGGAGGACCGAAGGAGCT
AACCGCTTTTTTGCACAACATGGGGGATCATGTAACTCGCCTTGATCGTTGGGAACCGGAGCTGAATGAAGCCATACCAAACGACGAGC
GTGACACCACGATGCCTGTAGCAATGGCAACAACGTTGCGCAAACTATTAACTGGCGAACTACTTACTCTAGCTTCCCGGCAACAATTA
ATAGACTGGATGGAGGCGGATAAAGTTGCAGGACCACTTCTGCGCTCGGCCCTTCCGGCTGGCTGGTTTATTGCTGATAAATCTGGAGC
CGGTGAGCGTGGGTCTCGCGGTATCATTGCAGCACTGGGGCCAGATGGTAAGCCCTCCCGTATCGTAGTTATCTACACGACGGGGAGTC
AGGCAACTATGGATGAACGAAATAGACAGATCGCTGAGATAGGTGCCTCACTGATTAAGCATTGGTAACTGTCAGACCAAGTTTACTCA
TATATACTTTAGATTGATTTAAAACTTCATTTTTAATTTAAAAGGATCTAGGTGAAGATCCTTTTTGATAATCTCATGACCAAAATCCC
TTAACGTGAGTTTTCGTTCCACTGAGCGTCAGACCCCGTAGAAAAGATCAAAGGATCTTCTTGAGATCCTTTTTTTCTGCGCGTAATCT
GCTGCTTGCAAAC

#### gBlock template sequence for *ACTB* insertions with ACTB_peg1 + ACTB_peg2 and ACTB+peg1 + ACTB_peg3

GCCACCATGGTGAGCAAGGGCGAGGAGCTGTTCACCGGGGTGGTGCCCATCCTGGTCGAGCTGGACGGCGACGTAAACGGCCACAAGTT
CAGCGTGTCCGGCGAGGGCGAGGGCGATGCCACCTACGGCAAGCTGACCCTGAAGTTCATCTGCACCACCGGCAAGCTGCCCGTGCCCT
GGCCCACCCTCGTGACCACCCTGACCTACGGCGTGCAGTGCTTCAGCCGCTACCCCGACCACATGAAGCAGCACGACTTCTTCAAGTCC
GCCATGCCCGAAGGCTACGTCCAGGAGCGCACCATCTTCTTCAAGGACGACGGCAACTACAAGACCCGCGCCGAGGTGAAGTTCGAGGG
CGACACCCTGGTGAACCGCATCGAGCTGAAGGGCATCGACTTCAAGGAGGACGGCAACATCCTGGGGCACAAGCTGGAGTACAACTACA
ACAGCCACAACGTCTATATCATGGCCGACAAGCAGAAGAACGGCATCAAGGTGAACTTCAAGATCCGCCACAACATCGAGGACGGCAGC
GTGCAGCTCGCCGACCACTACCAGCAGAACACCCCCATCGGCGACGGCCCCGTGCTGCTGCCCGACAACCACTACCTGAGCACCCAGTC
CGCCCTGAGCAAAGACCCCAACGAGAAGCGCGATCACATGGTCCTGCTGGAGTTCGTGACCGCCGCCGGGATCACTCTCGGCATGGACG
AGCTGTACAAGGGATCTGGTGCCACGAATTTCTCGCTACTCAAGCAGGCCGGTGATGTCGAGGAAAACCCTGGTCCTATGGATGATGAT
ATCGCCGCGCTCGTCGTCGACAACGGCTCCGGCATGTGCAAGGCCGGATTTGCAGGCGATGATGCTCCACG

**ssDNA donors used for 95nt insertion at *AAVS1***

ssDNA1: 5’ cggcATTACCCTGTTATCCCTAtcgGTACACCACTGAGACCGCGGTGGTTGACCAGACAT 3’
ssDNA2: 5’ cgaTAGGGATAACAGGGTAATgccgtctcggcatggacgagctgtacaaggaattctgag 3’

**Plasmid template sequences for 0.8/2.6k insertions at *TRAC***

0.8kb insertion template using the same template from AAVS1
2.4kb insertion template “pSLCAR-CD19-28z” (Addgene135991)
